# A Wheel of Fortune: the eukaryotic cell cycle as a tetra-stable excitable system

**DOI:** 10.1101/2025.08.31.673322

**Authors:** Calin-Mihai Dragoi

## Abstract

Progression through the eukaryotic cell cycle is governed by a complex biochemical network controlling the activation of cyclin-dependent kinases. Dynamically, the cell cycle control network can be viewed as a multi-stable system, in which nonlinear feedback loops generate multiple stable attractors, corresponding to distinct cell cycle phases (G1, S, G2, M). Transitions between these states are typically irreversible, ensuring ordered phase progression. However, recent studies of endocycles – variants in which subsets of phases are abrogated – have prompted the development of more general regulatory models, allowing for reversible transitions or the decoupling of autonomous oscillatory modules. The current paper combines minimal ODE modelling, phase plane analysis, and bifurcation theory to demonstrate that two prior frameworks – Newton’s Cradle and Latching Gate – are alternative representations of a shared architecture based on mutually regulated oscillators. Leveraging this insight, a generalized model is introduced that captures a broader spectrum of physiological endocycles than previously acknowledged. To provide an intuitive representation of the principles identified by the new model, a ‘Wheel of Fortune’ analogy for cell cycle dynamics is introduced.

## Introduction

The cell cycle is the sequence of events through which a growing eukaryotic cell replicates its components and partitions them between two daughter cells [1]. Central to this process is the accurate duplication and segregation of the genome, occurring in S phase (DNA synthesis) and M phase (mitosis), respectively. These phases are separated by gap phases (G1 and G2), and their ordered progression (G1–S–G2–M) is critical for genomic integrity. Disruptions—such as incomplete replication, DNA damage, or mitotic defects—trigger checkpoints that delay progression. This regulation is mediated by a protein network that modulate cyclin-dependent kinase (CDK) activity, with distinct CDK complexes driving S phase and mitosis.

Over the past three decades, experimental and theoretical studies have revealed that the CDK control network is structured around interlinked positive and double-negative feedback loops. These give rise to bistable switches and hysteresis, associating each cell cycle phase with a distinct stable steady state of the biochemical network [2,3]. Thus, biochemical processes that alter CDK activity dictate sharp and unidirectional transitions between phases, or checkpoint-mediated arrests. In this way, cells ensure faithful, ordered completion of essential physiological processes before returning to G1.

Extensively characterised variants of the cell cycle, known as endocycles [4,5], require an augmentation of this dynamical paradigm. In endocycles, cells abrogate one or more phases and periodically cycle through a reduced subset of events. In endoreplication, repeated G1–S transitions occur without mitosis, leading to successive doublings of chromosome numbers [4–6]. Similarly, in the natural process of meiosis [7] as well as under specific laboratory conditions [8–11], cells enter M-phase more than once without intervening DNA replication. In endomitosis, cells enter M-phase following G1/S/G2, but then abort the cycle prematurely [5,12,13]. This may result from a failure to divide or from perturbations of upstream cell cycle regulators [13]. In this paper, only the latter is considered.

Despite ample characterisation of cell cycle biochemistry, the dynamical principles governing endocycles remain hotly debated [8,11,14–17], thanks to their relevance in multicellular development, malignancy and cell cycle evolution. Several quantitative mechanistic models have been put forward [8,11,15], but inherent uncertainties in network connectivity and parameters, compounded by feedback and nonlinearity, resulted in inconsistent systems-level properties. Thus, no theoretical consensus exists presently.

Several dynamical frameworks stand out. One of the earliest proposals suggested that the CDK hysteresis oscillator is phase-locked with other oscillatory modules driving different events in the cell cycle [8,14]. Subsequently, the arrest of the CDK oscillator would allow these modules to run independently, with different phase/period, giving rise to endocycles [8]. The Latching Gate model [11,15] stands in counterpoint to this framework. Its assumption is that CDK activity is governed by an irreversible bistable switch, such that the two transitions controlling cell cycle entry and exit must be driven by distinct ‘helper molecules’. Endocycles occur when the switch is rendered reversible, such that one helper is sufficient to trigger both the ON and OFF transitions. The failure to engage the second helper abrogates half of the phases. Finally, in the Newton’s Cradle model [16], the CDK network comprises two mutually inhibitory oscillators, driving replication and mitosis. In normal mitotic cycles, the two oscillators alternate, as the activation of one oscillator induces a SNIC bifurcation and arrest of the other. Endocycles emerge when one oscillator is disabled and the other runs freely.

Although these models provide some conceptual clarity, they are limited in scope. So far, only replicative (G1/S) and mitotic (G2/M) endocycles are explained. Endomitosis – triggered by inhibition of specific mitotic regulators [13] - cannot be accounted for. Similarly, only two checkpoints can be explained without additional *ad hoc* mechanistic assumptions, despite the presence of at least three major checkpoints in most cells (G1/S, DNA damage, spindle assembly) [18].

To address these limitations, the present work develops dynamical models free from biochemical assumptions and identifies general structural properties of the phase space of endo-oscillatory systems. It is found that the Latching Gate and Newton’s Cradle share a common double-oscillator architecture. This insight is generalised into a dynamical model of cell cycle progression governed by a network of four oscillatory modules. This ‘Wheel of Fortune’ model permits selective disengagement of any module, thereby reproducing all known endocycles and predicting additional ones. It offers a flexible and minimal framework for understanding cell cycle dynamics across physiological and pathological contexts in eukaryotes.

## Results

### A minimal latching gate system

Endocycles were previously explained in two mechanistic models of the cell cycle – in budding yeast [15] and mammals [16] – using the Latching Gate hypothesis. Based on the yeast model, it was argued endocycles do not arise by the arrest of an oscillator in an entrained pair (cf. Lu & Cross, 2010 [8]), but through reversible switching of a hysteresis loop by helper molecules. This reasoning was later extended to the mammalian model, though certain phase plane and bifurcation properties differed between the two, leaving it unclear whether both systems shared the same underlying mechanism. Subsequently, the Newton’s Cradle model was introduced to explain the mammalian system as a pair of mutually inhibitory oscillators [16]. Here, the Latching Gate model is revisited in a minimal form to examine whether it represents a distinct endo-oscillatory mechanism or an alternative framing of Newton’s Cradle.

To this end, the premise of the Latching Gate hypothesis is assumed, implying the existence of a bistable species, A, engaged in negative feedback through two helper molecules, H1 and H2 (Fig. 1A). A inhibits H1 and stimulates H2. In turn, H1 activates A and H2 represses it.

**Fig 1.**
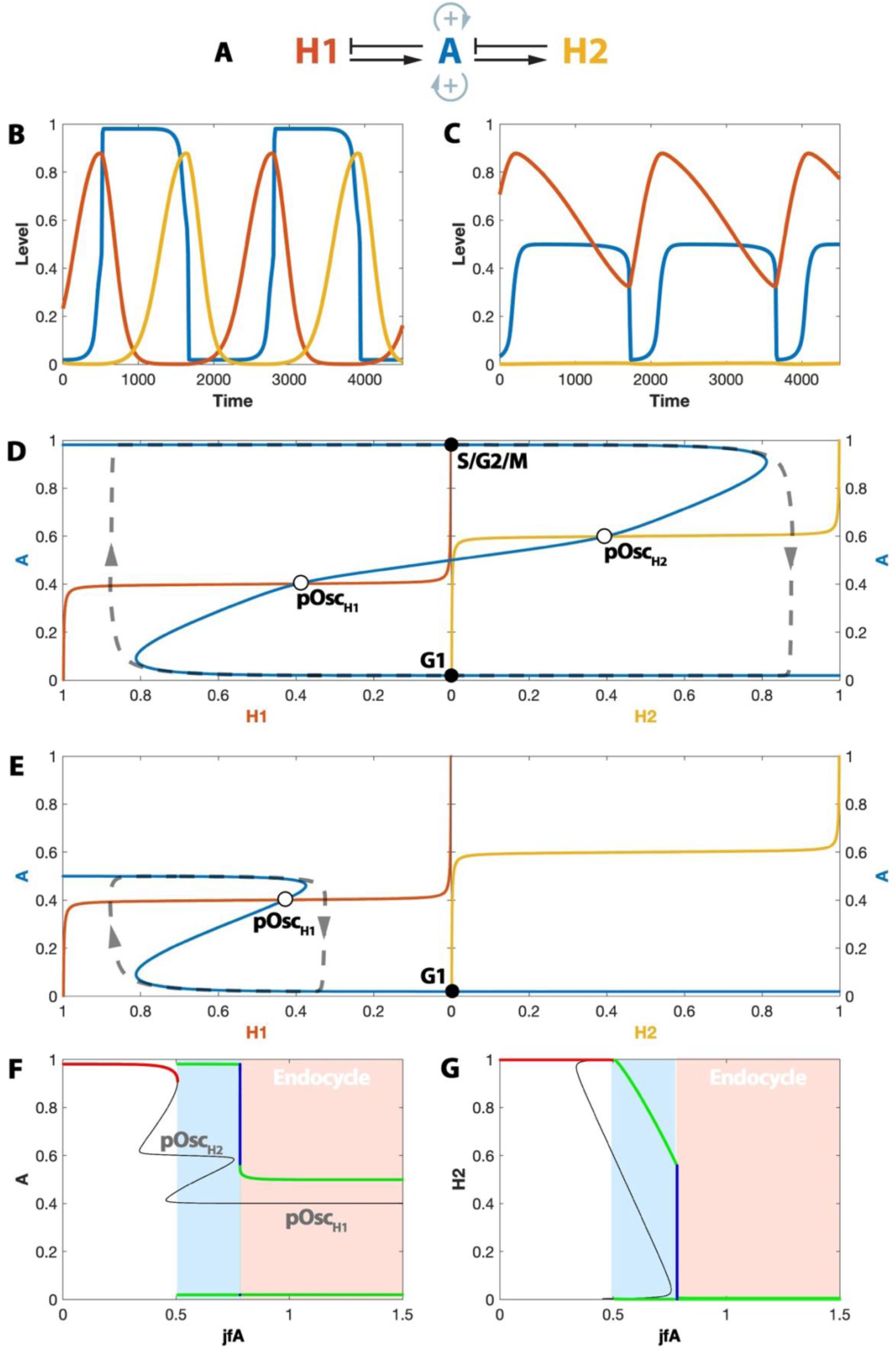
The dynamics of a minimal Latching Gates model. (A) Influence diagram for the A-H1-H2 system. (B) Time course simulation of the complete system oscillation. (C) Time course simulation of the H1 endocycle, with𝑗_𝑓𝐴_ = 1.2. (D) Representative sections of the A-H1-H2 phase portrait. Each half of the plot was calculated by computing the steady state solutions of one variable as a function of the other. The A-H1 phase plane (left-hand side) assumes H2=0. Similarly, the A-H2 phase plane (right-hand side) assumes H1=0. The stable steady states (within one particular plane, under the strict assumption that one of the H species is constant) are marked with black dots. The unstable steady states are marked with white dots. The time evolution of the system through these planes is marked with a grey dashed line. (E) The A-H1 and A-H2 phase planes in the H1 endocycle, with 𝑗_𝑓𝐴_ = 1.2. (F) Bifurcation diagram of A with respect to the parameter jfA. Stable steady states are marked in red, while unstable steady states are marked in black. The pOsc_H1_ and pOsc_H2_ unstable steady states are indicated. Green indicates the maximal and minimal amplitude of stable limit cycle solutions, while blue indicates the amplitude of unstable limit cycles. (G) Bifurcation diagram of H1 with respect to the parameter 𝑗_𝑓𝐴_ .

Thus, A is modelled as an ordinary differential equation, which assumes conversion between an active form A and an inactive one A’. Given a conservation law A + A’ = 1, A’ is not explicitly part of the model. Symmetrically, the forward and reverse conversion rates are characterised by nonlinear positive feedback, denoted by the functions B and B’:

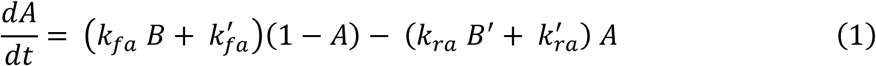

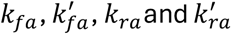 are dimensionless rate constants. B and B’ are Hill functions of A and (1 – A), respectively, and they depend on the helper molecules, H1 and H2, as follows:

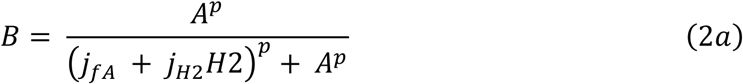

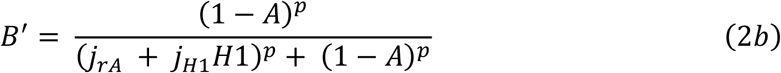

The threshold of these functions is a sum of a constant, 𝑗_𝑓𝐴_ (or 𝑗_𝑟𝐴_, respectively), and either H2 or H1 multiplied by a scaling factor (𝑗_𝐻2_or 𝑗_𝐻1_). The Hill exponent is denoted by *p*. These nonlinear positive terms allow for bistability in A [19].

The helpers in the original Latching Gate model had sigmoid shape. For this reason, H1 is modelled a symmetric conversion between H1 and H1’ = 1 – H1. The forward and reverse terms include a Michaelian-type function of H1 (1 – H1, respectively). This ensures that the steady state response of H1 is sigmoid with respect to A, for appropriate parameter choices. This is a well-established property of the Goldbeter – Koshland equilibrium of reversible, saturable processes [20].

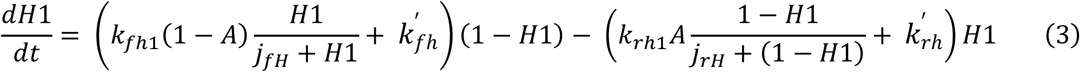

H2 is modelled analogously, but notice that the position of A and 1 – A has been swapped:

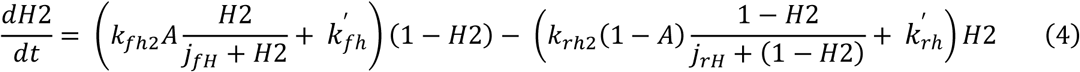

Although these equations are not a direct representation of any particular biochemical mechanism, they can be linked indirectly to cell cycle regulatory properties as outlined in Appendix A.

Given the parameter values in Table 1, numerical integration of this ODE system (Fig. 1B) reveals oscillations. Thanks to strong positive feedback, A alternates between high and low states, with ON and OFF transitions driven by H1 and H2, respectively. The system can switch to endocycles with a parameter change. Increasing 𝑗_𝑓𝐴_ from 0.6 to 1.2 (Fig. 1C) leads to an H1-driven endocycle, where H2 does not oscillate.

**Table 1.** Numerical values of parameters in the two-state endo-oscillatory system.

| Parameter | Description | Value |
| --- | --- | --- |
| kfa | Activation rate of A (autocatalytic term) | 0.5 |
| kra | Inactivation rate of A (autoinhibitory term) | 0.5 |
| kfa' | Basal activation rate of A | 0.01 |
| kra' | Basal inactivation rate of A | 0.01 |
| jfA | Activation threshold for A | 0.6 |
| jrA | Inhibition threshold for A | 0.6 |
| jH1 | Strength of H1-mediated inhibition of A | 0.5 |
| jH2 | Strength of H2-mediated activation of A | 0.5 |
| p | Hill exponent of A autocatalytic terms | 15 |
| kfh1 | Activation rate of H1 (autocatalytic term) | 0.4 |
| krh1 | Inactivation rate of H1 (autoinhibitory term) | 0.6 |
| kfh2 | Activation rate of H2 (autocatalytic term) | 0.4 |
| krh2 | Reverse inactivation rate of H2 (autoinhibitory term) | 0.6 |
| kfh' | Basal activation rate of H1 and H2 | 0.00001 |
| krh' | Basal inactivation rate of H1 and H2 | 0.00001 |
| jfH | Activation threshold for H1 and H2 | 50 |
| jrH | Inhibition threshold for H1 and H2 | 50 |

These oscillations are analysed using phase planes. Although the system is three-dimensional, it can be understood by examining two 2D cross-sections, since H1 and H2 are mutually exclusive (Fig. 1B). Each panel of Fig. 1D shows the nullclines (steady-state curves) of A and one helper, with the other helper set to zero. The variable A functions as an irreversible bistable switch in both cases: the ON transition (via saddle-node bifurcation) occurs only when H1 is active, and the OFF transition only when H2 is active. The nullcline of A intersects the sigmoidal nullcline of H1 at three points. The intersection on the upper branch is stable and labelled S/G2/M (cf. Novak and Tyson, 2022). Two other intersections lie on the unstable branch of A, one labelled pOsc_H1_ (pseudo-oscillator) and another left unlabelled. The right panel shows the anti-symmetric case for H2.

This view explains the time evolution: starting at G1, H1 accumulates until the lower branch of A loses stability and A switches on. Once A=1, H1 is no longer stimulated, locking the system in S/G2/M. To return to G1, the system transitions similarly via the A– H2 plane. When 𝑗_𝑓𝐴_ is increased (Fig. 1E), the upper branch no longer crosses the A–H2 plane, rendering the cycle reversible with respect to H1 and thus driving an endocycle.

A bifurcation diagram with respect to 𝑗_𝑓𝐴_ (Fig. 1F) reveals how the system transitions from the normal cycle to the endocycle regime, as pOsc_H2_ merges with an unstable steady state. Concurrently, the amplitude of oscillation sharply drops to zero for H2 (Fig. 1G), but remains robust for H1 (Fig. S1A). The oscillation period also lengthens dramatically, suggesting critical slowing down, consistent with a global (SNIC) bifurcation (Fig. S1B), as explored further below.

The pseudo phase planes in Fig. 1 match those of the original Latching Gate model (Novak & Tyson 2022, Fig. 3) but differ from those of Newton’s Cradle (Dragoi et al. 2024, Figs. 6 and 9). This confirms the two systems are both endo-oscillatory without being dynamically identical. However, they share many essential features, as shown next.

### The latching gate endo-oscillatory system comprises two oscillators

Previous work [11,15] and Fig. 1 argue that endocycles emerge when one of the stable steady states (G1 or S/G2/M) is lost, rendering the switch reversible with respect to one of the helpers. However, Fig. S1A shows exceptions. At 𝑗_𝑓𝐴_ = 0.8 – a value within the endocycle region of the bifurcation diagrams (Fig. 1F,G; Fig. S1B) – an H1 endocycle arises, as in Fig. 1E. Yet, the upper branch of A still appears in the A–H2 plane, and S/G2/M remains intact. Instead, an intermediate branch of the A nullcline prevents access to S/G2/M.

Because of this, the A nullcline alone cannot be used as a reliable diagnostic of endo- oscillatory behaviour. An alternative indicator lies in the status of the unstable steady state pOsc_H2_, which is absent both at 𝑗_𝑓𝐴_ = 0.8 (Fig. S1A) and 𝑗_𝑓𝐴_ = 1.2 (Fig. 1E). This suggests that the A–H1–H2 system contains two nonlinear oscillators and that endocycles result from the disappearance of one. In Fig. 1, the transition to endocycles coincides with the loss of pOsc_H2_. This behaviour is consistent with the Newton’s Cradle model [16], where endo-oscillations are driven by two amplified negative feedback loops (two activators and two inhibitors). Yet, here, only three species are present, raising the question of how two oscillators can emerge.

The explanation lies in the two positive feedback loops A engages in. Due to high nonlinearity (Hill exponent = 15), the two feedback motifs predominate at high and low A, respectively. This is because the activation thresholds are set by H1 and H2, which are turned on and off by A. This can be visualised more clearly by imagining A as split into two components – A_High_ and A_Low_ – as shown in Fig. 2A. Under this framing, the A–H1–H2 system appears as a pair of coupled oscillators. To test this interpretation, the system was perturbed to eliminate one pOsc state (specifically, pOsc_H2_) without directly modifying the A nullcline. The H2 nullcline is sigmoidal, with activation threshold controlled by 𝑘_𝑓ℎ2_ and 𝑘_𝑟ℎ2_. Assuming 𝑘_𝑟ℎ2_ = 1 – 𝑘_𝑓ℎ2_, the threshold equals 𝑘_𝑓ℎ2_. Reducing 𝑘_𝑓ℎ2_ lowers this threshold, eventually causing pOsc_H2_ to vanish via a homoclinic bifurcation, as it merges with a nearby unstable steady state (Fig. 2B; 𝑘_𝑓ℎ2_ = 0.7). The resulting endocycle is driven by pOsc_H1_. A oscillates with reduced amplitude and H2 remains close to 1 (Fig. 2C). As in other endocycles, the transition is abrupt (Fig. 2D) and displays critical slowing down (not shown).

**Fig. 2.**
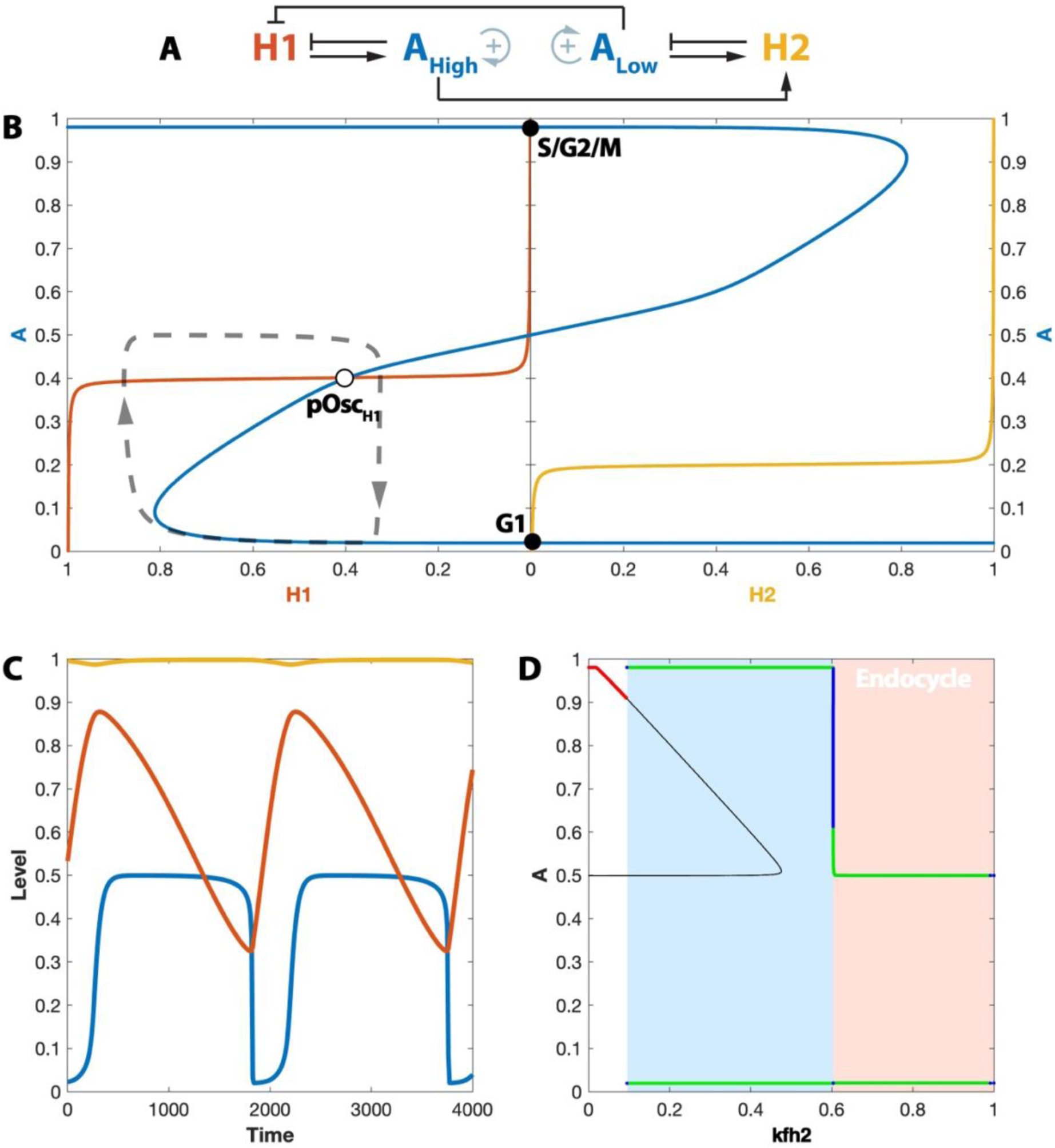
An alternative mechanism for endocycle induction. (A) Influence diagram for the A-H1-H2 system, highlighting its double-oscillator organisation. (B) The A-H1 and A-H2 phase planes in the H1 endocycle, with 𝑘_𝑓ℎ2_= 0.7. The A-H1 phase plane (left-hand side) assumes H2=0. Similarly, the A-H2 phase plane (right-hand side) assumes H1=0. (C) Time course simulation of the H1 endocycle, with 𝑘_𝑓ℎ2_=0.7. (D) Bifurcation diagram of A with respect to 𝑘_𝑓ℎ2_.

Because H2 ≈ 1 in this regime, the standard phase planes – assuming H2 = 0 (left) or H1 = 0 (right) – become misleading. A new set of pseudo-phase planes with H2 = 1 (left) and H1 = 1 (right) is plotted in Fig. S1D. Under these conditions, A forms a reversible bistable switch with both helpers. On the A–H1 plane, this produces pOsc_H1_, which drives the endocycle. On the A–H2 plane, a new stable steady state at (1, 0.5) emerges, fixing H2 at high levels.

These observations support the view that the pOsc attractors act as autonomous limit cycle oscillators. When both are present, the system performs full oscillations: A cycles through its full range, and both helpers engage in turn. In this case, each pOsc behaves as a ‘pseudo-oscillator’, temporarily inactive while the other operates. To generalise this framework beyond endoreplication and mitotic endocycles (H1 and H2 endocycles), a mechanism for coordinating multiple pseudo-oscillators must be identified. For this, the A–H1–H2 system is next analysed from a different perspective.

### The Latching Gate model as a bistable excitable system

Generalising the double phase-plane analysis to systems with more than two pseudo- oscillators is not straightforward. To address this, variable A is reformulated as two coupled ODEs, enabling analysis of how it transitions between its two stable steady states under the influence of helper variables. Specifically, function B is redefined as a dynamic variable whose steady state matches its original definition:

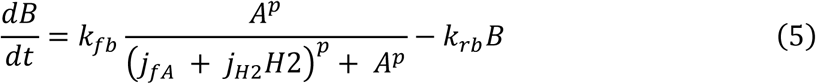

Here 𝑘_𝑓𝑏_ =𝑘_𝑟𝑏_, and both are set much larger than all other rate constants in the model, ensuring B remains in pseudo-steady state relative to the rest of the system.

The A–B phase plane can then be plotted with H1 = H2 = 0 (Fig. 3A). The A-nullcline (dark blue) is bistable with respect to B, while the B-nullcline (light blue) is sigmoidal, as expected from B’s Hill-type steady state function. Their intersections yield two stable steady states - G1 (low A) and S/G2/M (high A) – as well as one intermediate unstable state (unlabelled). A representative trajectory is overlaid with dashed/dotted lines and colour coded.

**Fig. 3.**
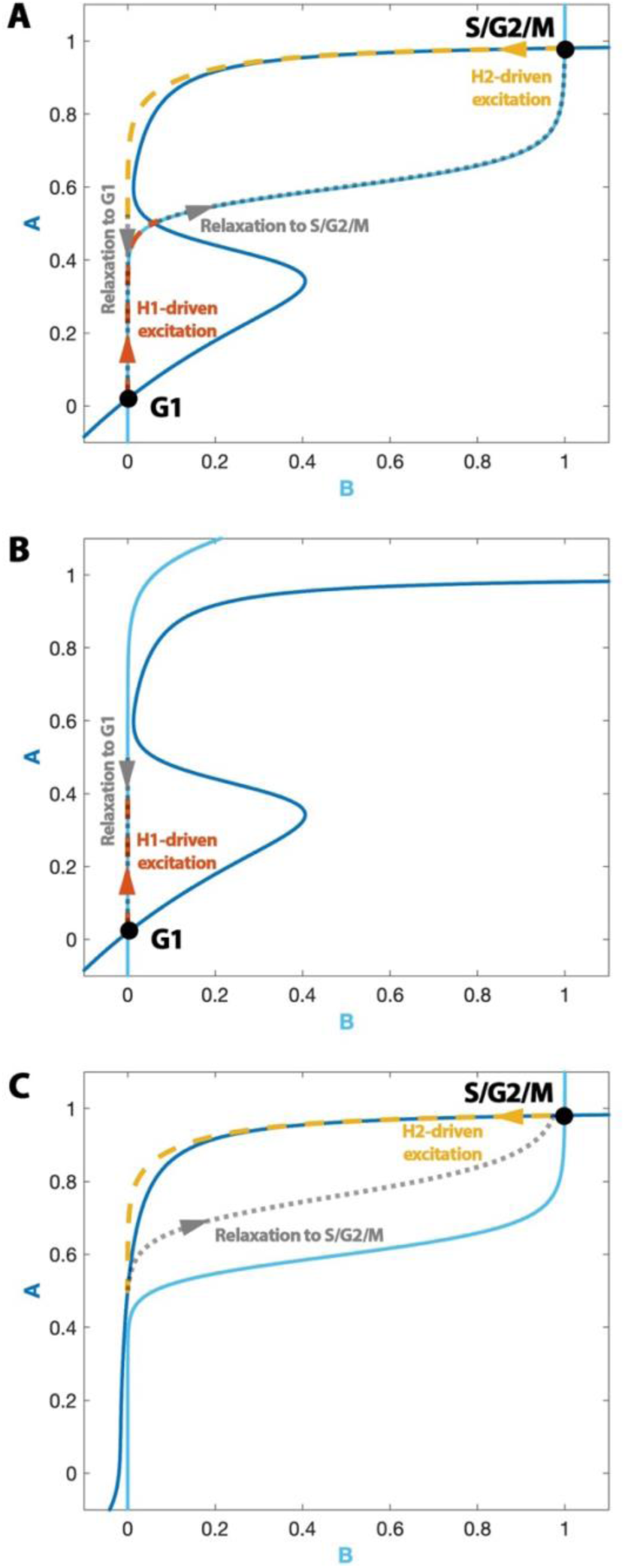
The latching gates model from the core bistable switch perspective. (A) Phase plane for the AB variables, assuming H1=H2=0. In the absence of helpers, the phase portrait shows two stable steady states, corresponding to the G1 and S/G2/M, consistently with Fig. 1D. The simulated trajectory of the system is plotted in dotted/dashed lines. Orange and yellow indicate the system dynamics dependent on the action of H1 and H2, as required to escape the basins of attraction of G1 and S/G2/M, respectively. Dotted grey lines show spontaneous relaxations of the A-B system to one of the stable steady states. (B) AB phase plane with 𝑗_𝑓𝐴_= 1.2, showing the loss of the S/G2/M state and the emergence of the H1 endocycle. (C) AB phase plane with 𝑗_𝑟𝐴_= 1.2, showing the loss of the G1 state and the emergence of the H2 endocycle.

Starting at G1, H1 accumulates, pushing A up along the B-nullcline near A ≈ 0.5. From Fig. 1C (H1 endocycle), it is known that H1 alone cannot drive A accumulation past this threshold. Once A reaches this intermediate level, the system is spontaneously attracted to the S/G2/M state and relaxes along the B-nullcline. Now, H2 accumulation begins, reducing both A and B. Once A is sufficiently suppressed, the system returns to G1.

This phase-plane perspective offers a new way to understand endocycles. As shown previously (Fig. 1C,E), increasing 𝑗_𝑓𝐴_ to 1.2 leads to H1-driven endocycles. On the A–B plane, this perturbation shifts the B-nullcline to the right (Fig. 3B), eliminating the S/G2/M state via a saddle-node bifurcation. Thus, after excitation by H1, the system has no state to relax to other than G1 – producing an ‘abridged’ oscillation. The H2 endocycle operates analogously (Fig. 3C). As per the previous section, the loss of G1 or S/G2/M is a sufficient but not necessary condition for endo-oscillations. However, this formulation establishes a foundation for generalising the latching gate framework to systems with more than two coordinated pseudo-oscillators.

### A tetra-stable core system

The phase-plane framework introduced above can be extended to systems with more pseudo-steady states (i.e. cell cycle phases) by reshaping the nullclines of variables A and B and introducing appropriate helper molecules. Suppose that both A and B are subject to nonlinear positive feedback (Fig. 4A), according to the equations in Materials and methods and parameters in Table 2. In this case, each variable is bistable with respect to the other. Their nullclines can then intersect at four points on their stable branches, yielding four stable steady states (Fig. 4B) labelled G1, S, G2, and Metaphase (naming explained in the Discussion). The intersections of stable and unstable branches, correspond to saddle points, which lie on the boundary of a stable state’s basin of attraction (Fig. S2). These steady states can be seen as ‘thresholds’ for cell cycle phase transitions, so they are labelled Th_G1/S_, Th_S/G2_, Th_G2/Meta_, Th_Meta/G1_. Importantly, a negative feedback loop is assumed, with A activating B and B inhibiting A. This interaction generates a fifth steady state at the intersection of the nullclines’ unstable branches, termed the pseudo-oscillator (pOsc). This key design feature is revisited below and in the Discussion.

**Fig. 4.**
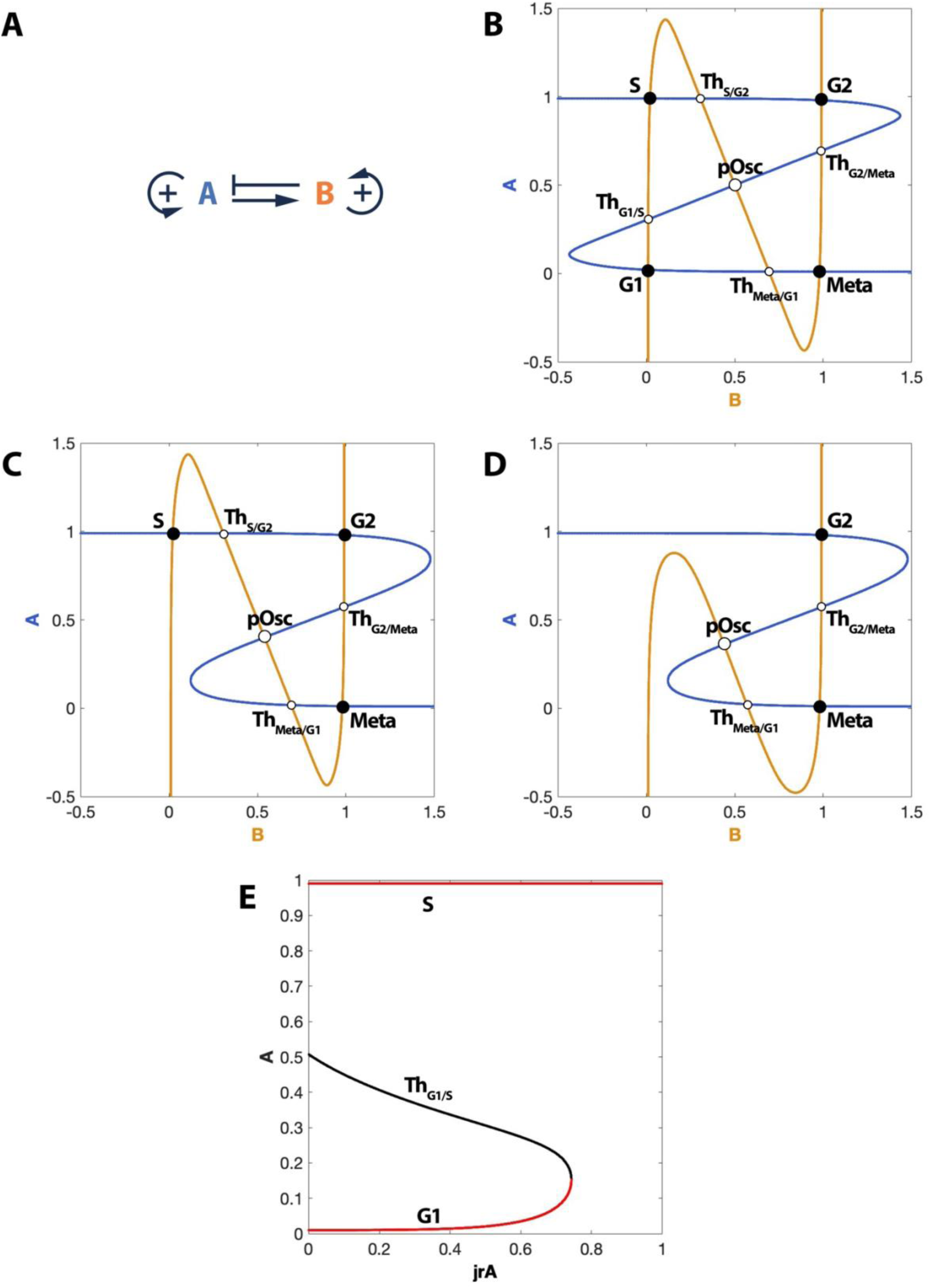
A tetra-stable core system. (A) Influence diagram of the core tetra-stable system. (B) Phase plane for the basal AB core system. (C) Phase plane for the core system with 𝑗_𝑟𝐴_=0.8, where the G1 state is eliminated. (D) Phase plane for the core system with 𝑗_𝑟𝐴_=0.8 and 𝑗_𝑟𝐵_=0.8, where the G1 and S states are eliminated. (E) Bifurcation diagram of A with respect to the 𝑗_𝑟𝐴_ parameter. Bifurcation diagram legend: red denotes stable steady states and black denotes saddle and unstable points.

**Table 2.**
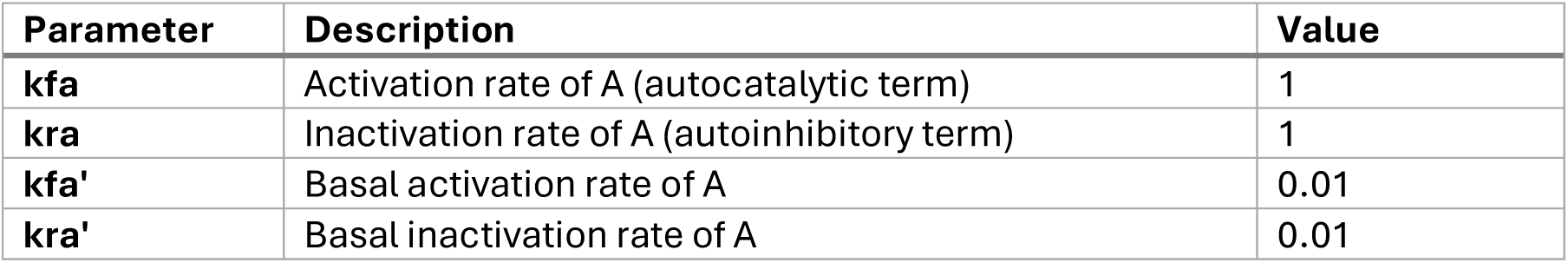
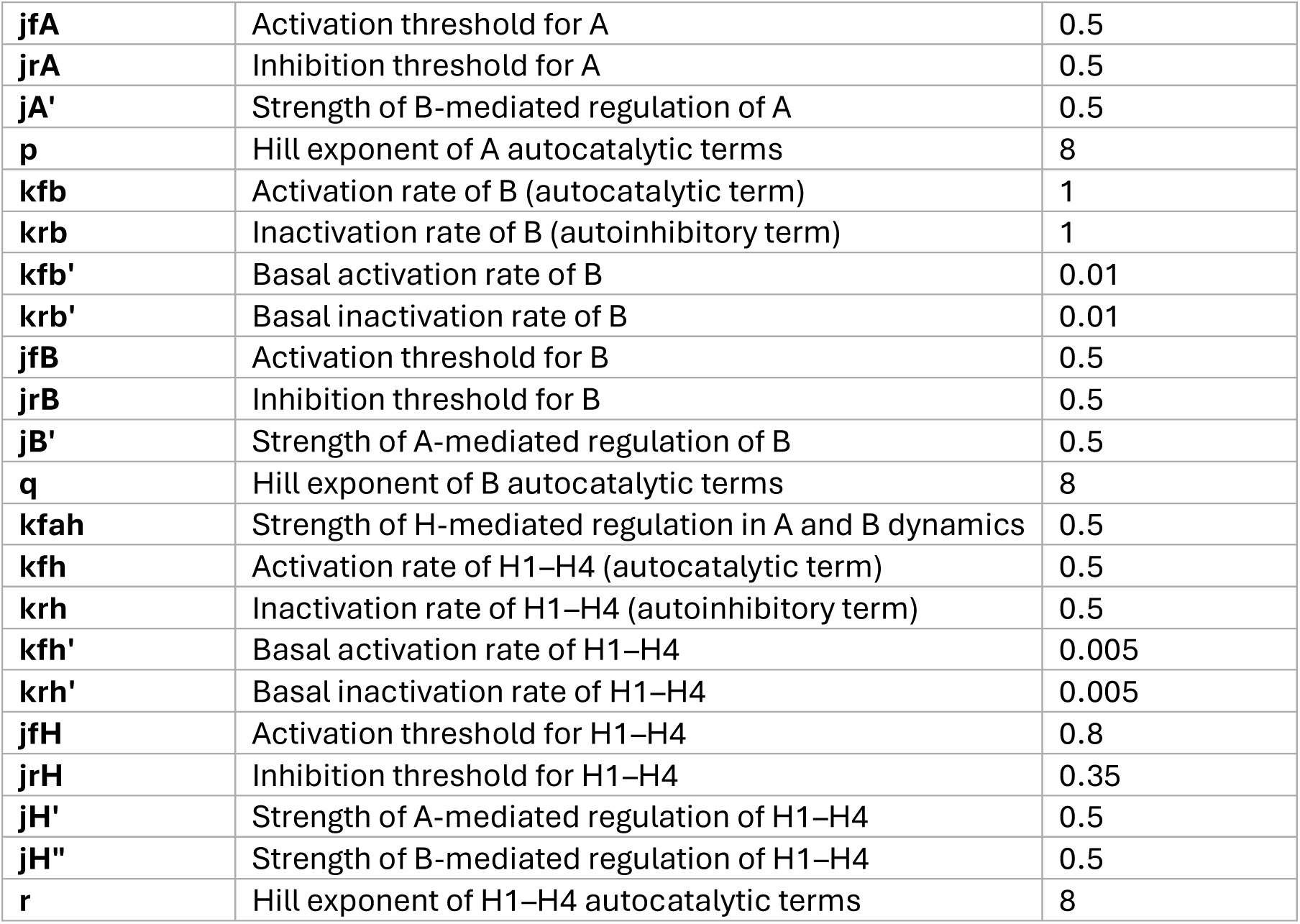
Numerical values of parameters in the four-state endo-oscillatory system.

The tetra-stable core could support periodic oscillations if each steady state triggered the activation of a helper molecule that, in turn, excites the system across the corresponding phase threshold. For example, starting from G1, the activation of a helper (H1) could push the system into the S state by increasing A. Such an excitation process can be driven by a helper–core pseudo-oscillator, as detailed in the next section.

In this system, an endocycle arises when one or more stable steady states are eliminated. For instance, increasing the parameter 𝑗_𝑟𝐴_ shifts the activation threshold of A’s nullcline to higher values (Fig. 4C). This perturbation removes the G1 steady state, as it merges with Th_G1/S_, at a saddle-node bifurcation (Fig. 4E), causing the system to transition directly to S. In this scenario, G1 is skipped, and the system progresses to the next phase without the need of H1. This is consistent with the interpretation of an endo- oscillation as a limit cycle in which one pseudo-oscillator is disengaged. To prevent the engagement of the helper, its dynamics must evolve on a timescale much slower than those of A and B. As a result, the system passes quickly through the former G1 region, spending insufficient time at low A and B to support a full G1 program.

Thanks to symmetries in the A–B system, similar bifurcations can eliminate any subset of the four steady states. This flexibility allows the tetra-stable core to support up to 15 distinct endocycles (whether physiological or not). For example, eliminating both G1 and S (Fig. 4D) produces a mitotic endocycle in which the system passes quickly through early phases and is not expected to support DNA replication.

The negative feedback between A and B plays a crucial role in maintaining directionality during endocycles. If all four stable steady states (G1, S, G2, and Meta) are suppressed, the resulting pOsc state drives a clockwise limit cycle oscillation in the A–B phase plane. When only some phases are eliminated, this oscillatory core ensures that the system continues progressing in the correct order, even in the absence of the corresponding helper activity. Metaphorically, the central negative feedback acts as a form of dynamical inertia, sustaining the system’s trajectory when helper-induced ‘forces’ are unavailable. In contrast, the simpler bistable model discussed earlier does not require negative feedback between A and the pseudo-species B, as it involves excitations and relaxations around a single pseudo-steady state, without the need for ordered progression through multiple phases.

### The excitatory module: helper-driven negative feedback pseudo-oscillators

As in the bistable model, helper molecules drive transitions between states in the core module. For this to occur in the correct sequence, each helper must be activated only when the system is in its corresponding phase. For example, the G1/S helper (H1) should only become active in the G1 state, while the S/G2 helper (H2) activates in S, and so on. Since each core state is defined by a unique combination of A and B activities, these variables can regulate helper expression. In particular, G1 is characterized by low A and low B. Therefore, H1 must be suppressed in any state where either A or B is high, implying that both A and B inhibit H1 (Fig. 5A). Conversely, since S differs from G1 primarily in having high A, H1 promotes the G1→S transition by activating A.

**Fig. 5.**
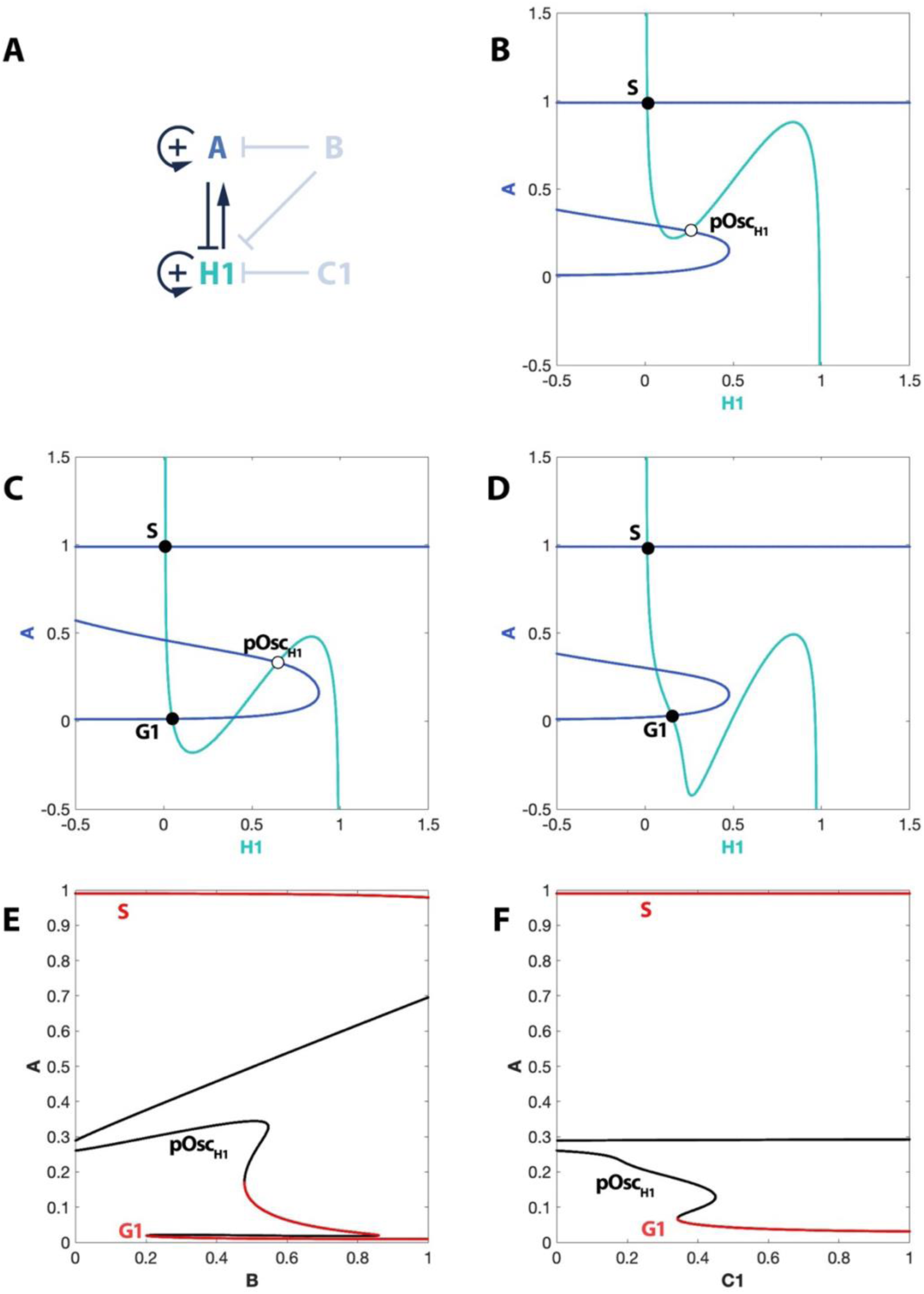
The dynamical properties of the excitatory module. (A) Influence diagram for the excitatory module associated with the G1 state. H1 and A are dynamical variables here, while B and C1 (light grey) are taken as parameters. (B) Phase plane of the excitatory module with B=0. (C) Phase plane with B=0.4. (D) Phase plane with the checkpoint parameter C1=1. (E) Bifurcation diagram of A with respect to the parameter B. (F) Bifurcation diagram of A with respect to the checkpoint parameter C1.

To analyse this interaction, the H1–A excitatory module is considered, with B held constant as a parameter. Due to system symmetry, all helper modules behave analogously. In the H1–A phase plane (Fig. 5B, B=0), the A nullcline is bistable due to nonlinear positive feedback, consistent with the bistability of A in the full core system (Fig. 4B). Because H1 activates A, the high-A branch becomes the only stable outcome when H1 is large. Meanwhile, H1 also self-activates, rendering its nullcline bistable as well. The negative feedback between A and H1 gives rise to an unstable steady state at the intersection of their unstable branches, denoted pOsc_H1_. While this configuration is capable of oscillation, in the presence of the stable S-state, the oscillation is blocked – so the helper simply acts as a driver, pushing the system over the Th_G1/S_ threshold into S phase.

A difference from the bistable model is that the helper (H1) is bistable, rather than merely sigmoid, due to auto-amplification. This ensures high oscillation amplitude for the helper. This property is of relevance, as in the current context, endocycles are explicitly defined in relation to the helper disengagement, as explained in the previous section, rather than the loss of the pOsc_H_ state, which was the key hallmark of endocycles in the bistable system. However, more importantly, this feature improves the tunability of the oscillator [21] and reflects known properties of cell-cycle regulators [16].

The excitatory module’s behaviour depends on the core state. When A is high (e.g., in the S phase), the system immediately reaches the S steady state on the H1–A plane, with little further H1 change. However, when B increases, it suppresses H1 by translating the H1 nullcline downward, allowing a new low-H1, low-A stable state to emerge, corresponding to G1 (Fig. 5C). This mechanism prevents premature H1 accumulation, particularly in the Meta state (high B, low A), where transition back to G1 must not immediately trigger H1 reactivation.

Perturbations that inhibit H1 activation mimic the effect of high B. For example, a checkpoint control parameter, C1, shifts the H1 nullcline down, suppressing H1 regardless of B activity (Fig. 5D). When C1 is high, the G1 steady state persists, and the system cannot proceed to S, thus arresting oscillation. Bifurcation diagrams of the H1–A module with respect to B and C1 (Figs. 5E, 5F) show that the G1 state appears via a saddle-node bifurcation, with thresholds near 0.2 and 0.35, respectively. This sharp onset of arrest mirrors prior models of checkpoint dynamics [22,23].

### The dynamics of the complete mitotic oscillation

To model the full cell cycle, the system must autonomously oscillate through all core module states in sequence. This is achieved by introducing helpers H2, H3, and H4, analogous to H1 (Fig. 6A). H2 accumulates in the S state (A high, B low), being activated by A and inhibited by B. In turn, H2 activates B, driving the S→G2 transition. In G2, both A and B are high, activating H3, which inactivates A to reach Metaphase. The H3–A module is anti-symmetric to the H1–A module. Finally, H4 activates B from low levels, completing the Meta→G1 transition; the H4–B module is anti-symmetric to H2–B. Time-course simulations (Fig. 6B) confirm the expected sequence: H1 and A rise in G1, followed by H2 and B in S and G2, then H3 removes A to reach Meta, and H4 resets the system to G1.

**Fig. 6.**
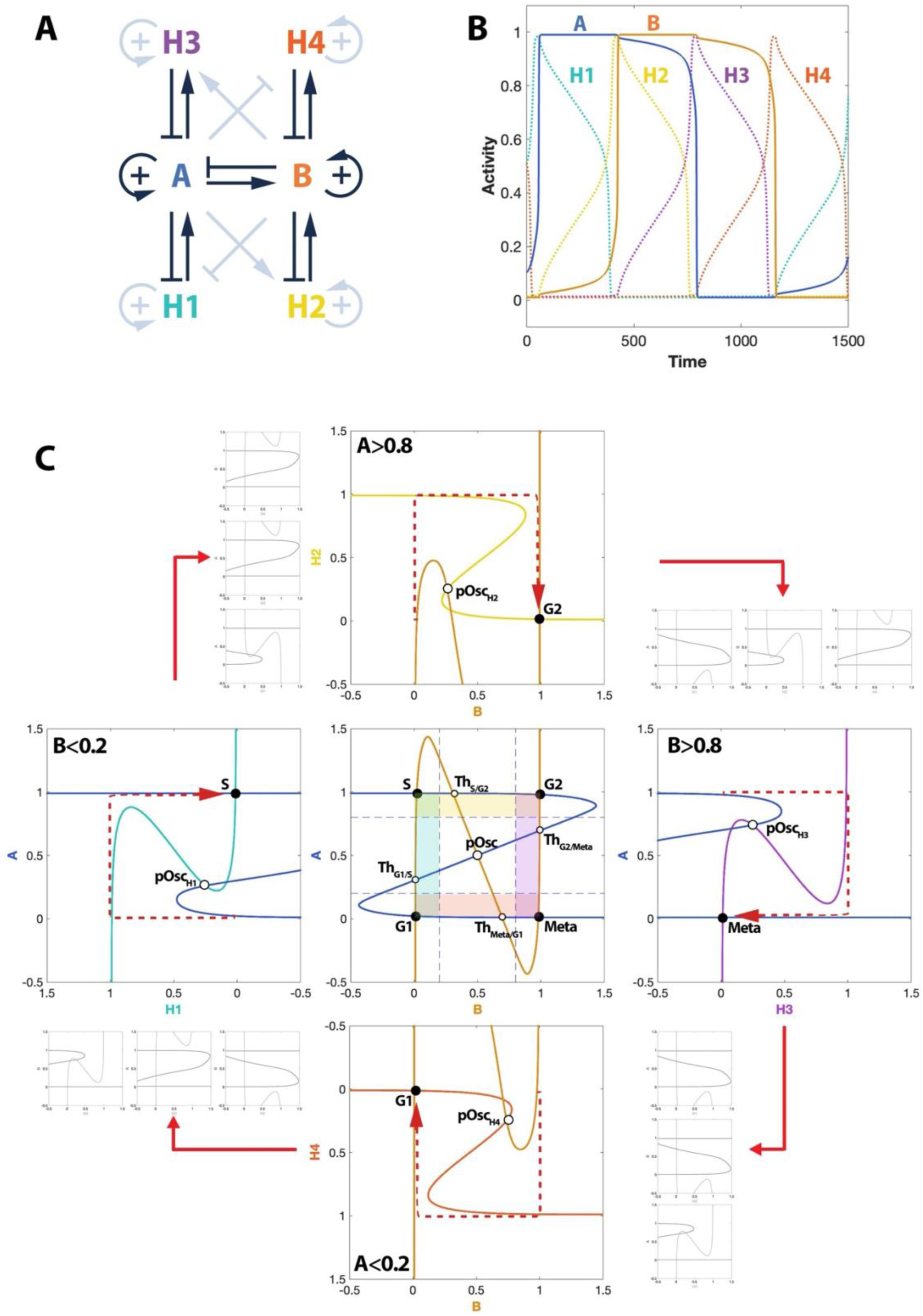
The dynamics of the complete mitotic oscillator. (A) Influence diagram. (B) Time course simulation. (C) The oscillation trajectory through projections of the 6-dimensional phase space. The central panel is the phase plane of the core model when all helpers are zero. The coloured regions represent the portions of the phase plane where each helper is active, bounded by the 0 and 1 values of each core variable. The phase planes on the periphery show each excitatory module at each phase of the core system. The enlarged, coloured phase planes correspond to the helpers driving the particular transitions, while the adjacent small grey panels show the state of the other excitatory modules at the same time. The dotted red arrows indicate the calculated trajectory on each helper’s phase plane.

The mechanism can be visualized using 2D phase-plane slices (Fig. 6C). In the A–B core plane (helpers off), each state appears trapped in its stable attractor. However, in the A– H1 plane (B=0), G1 is absent; the only stable state is S, so H1 accumulation carries the system over the G1→S barrier. In the B–H2 plane (A=1), S is absent, enabling S→G2, and so forth until G1 is reached again. Each helper module lies in a plane orthogonal to the core variables, effectively making the system ‘jump’ between attractor basins, over the barriers in the A–B plane.

An alternative view examines the core phase plane under active helpers (Fig. 7). With helpers off, the tetra-stable landscape of Fig. 4B applies. When H1=1 (G1/S transition), G1 disappears, and the system is attracted to S spontaneously. When H2 accumulates, S vanishes, driving the system to G2; the pattern repeats for H3 and H4. Of course, the trajectory of the system need not be confined to these 2D sections of the phase spaces, but this perspective provides a useful intuitive picture of how helpers reshape the core dynamics during transitions.

**Fig. 7.**
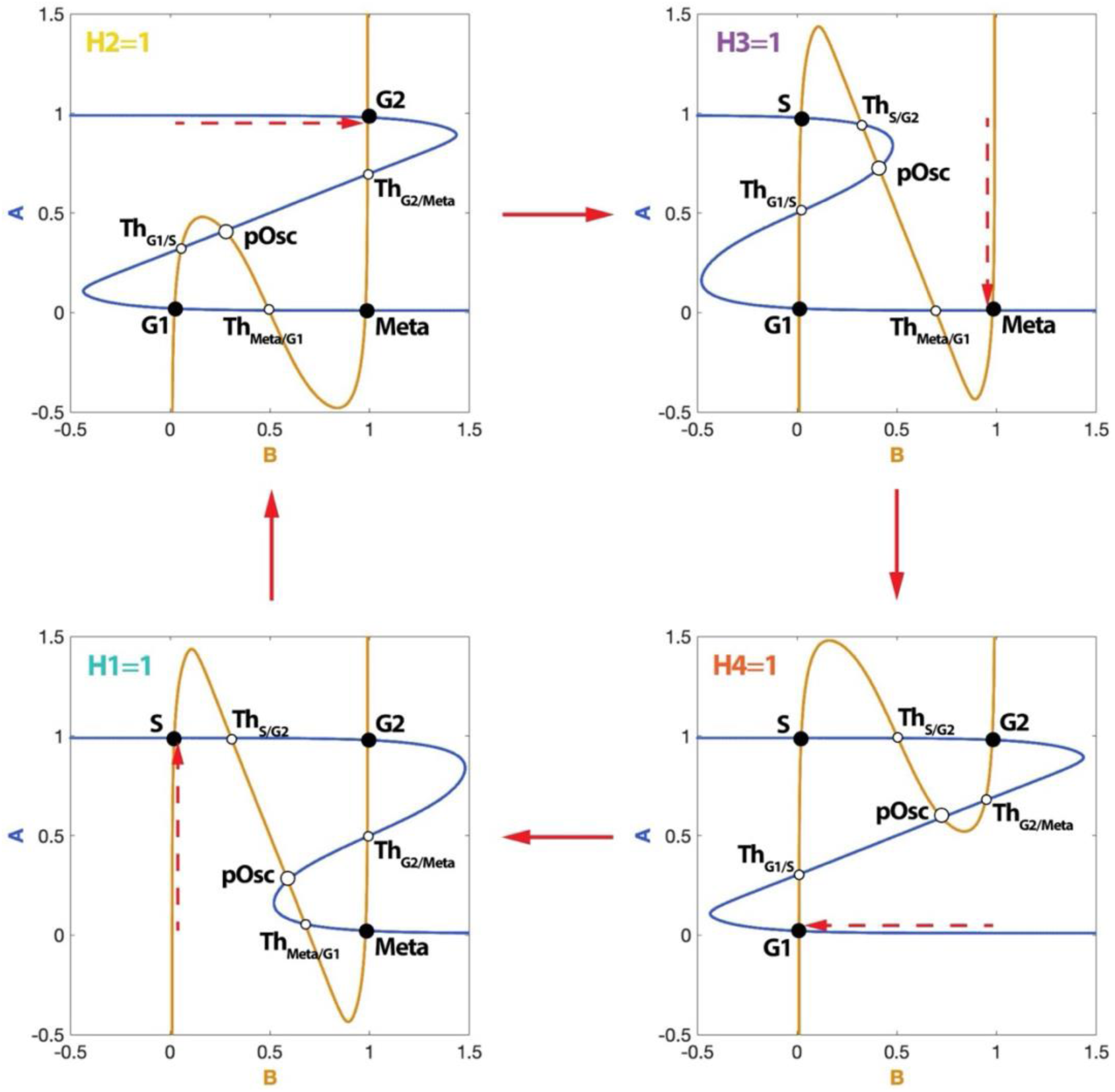
An alternative view of the oscillator’s dynamics. the plot shows the sequence of changes to the core phase plane as each helper is activated. When each helper is sufficiently active, the state that led to its activation loses stability via a saddle-node bifurcation and the system is attracted to the next stable steady state in line.

### Endocycle dynamics

Endocycles are here defined as perturbed periodic solutions in which a specific helper module is disengaged. In the core system (Fig. 4C,D), this occurs when one of the four stable steady states is removed via single-parameter changes to 𝑗_𝑟𝐴_, 𝑗_𝑓𝐴_ , 𝑗_𝑟𝐵_, or 𝑗_𝑓𝐵_ (see Methods), which shift the ON/OFF thresholds of the A and B nullclines. Eliminating a state removes the barrier for entry into the next state, allowing the core to bypass the corresponding helper. Since helpers evolve more slowly than the core, they cannot accumulate significantly in these conditions, and the targeted helper effectively drops out of the oscillation. For example, with 𝑗_𝑟𝐵_=1, the S state is suppressed and H2 barely rises (Fig. 8A). Suppressing two states yields more complex endocycles: with 𝑗_𝑓𝐵_ =𝑗_𝑟𝐵_=1, both S and Meta vanish, disengaging H2 and H4 (Fig. 8B).

**Fig. 8.**
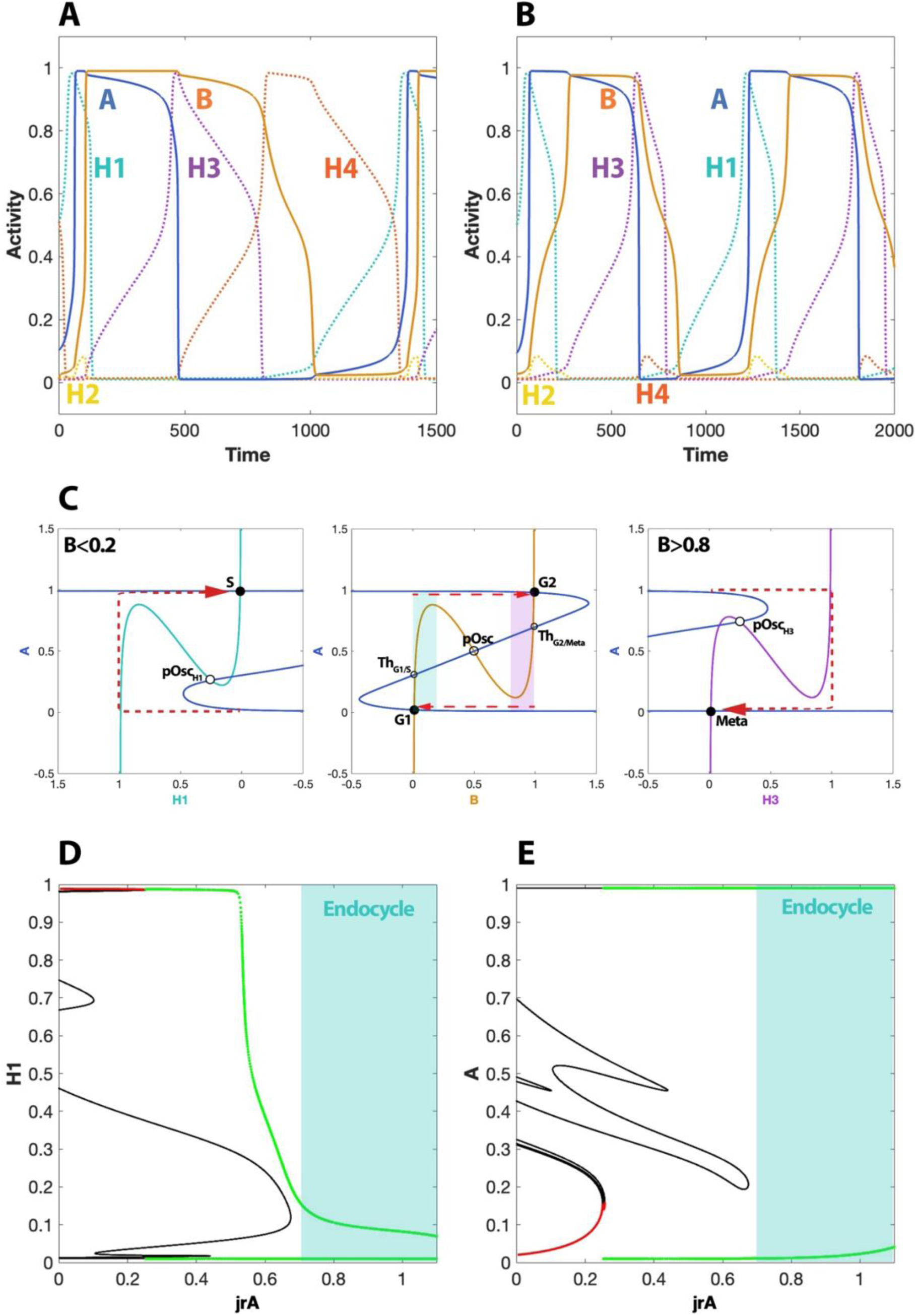
Endocycle dynamics. (A) Time course simulation for 𝑗_𝑟𝐵_=1, showing H2 disengagement from oscillation. (B) Time course simulation for 𝑗_𝑟𝐵_=𝑗_𝑓𝐵_=1, showing H2 and H4 disengagement from oscillation. (C) Phase plane view of the 𝑗_𝑟𝐵_=𝑗_𝑓𝐵_=1 perturbation. The G1/S and G2/Meta transitions are driven by excitatory modules (shown by left and right panels), while the S/G2 and Meta/G1 transitions are spontaneous for the core. (D) H1 vs 𝑗_𝑟𝐴_bifurcation diagram showing the dramatic loss of amplitude in H1 oscillation with 𝑗_𝑟𝐴_increase. (E) A vs 𝑗_𝑟𝐴_bifurcation diagram showing virtually no change in A amplitude with 𝑗_𝑟𝐴_increase. Bifurcation diagram legend: red denotes stable steady states, black denotes saddle and unstable points, and green denotes oscillation amplitude.

This endocycle can be visualised on the projections through the phase space (Fig. 8C), in analogy to Fig. 6C. The G1/S and G2/Meta transitions take place in the plane of the excitatory modules, under the action of H1 and H3, respectively. In contrast, the S/G2 and Meta/G1 occur spontaneously in the A–B core plane.

The transition from the mitotic cycle to endocycles can be characterised by plotting a bifurcation diagram of a helper variable against the parameter that controls the stability of its inducing state. In Fig. 8D, H1 maintains a high amplitude for 𝑗_𝑟𝐴_ ≲ 0.6; beyond this threshold, the amplitude drops abruptly to <15% of the unperturbed value, while other helpers remain unaffected (Fig. 8E). This threshold is taken as an operational definition of an endocycle in the tetra-stable system. This contrasts with the two-state system, where endocycles emerge through oscillator loss via a global bifurcation (Fig. 1F,G). In the tetra-stable case, the pOsc_H_ state persists, but the system transitions away from the helper’s activation phase too rapidly for significant accumulation.

### The latching gate perspective on the four-state system

The initial four-phase system was introduced in core A-B form because the latching gate view did not generalise easily beyond two phases. With a general four-state version now available, we can revisit this perspective. In the bistable case (Fig. 1D), a pair of phase planes of A versus each helper sufficed; here, four planes are required. For the tetra- stable system, the A–H1 and A–H3 planes are arranged analogously (Fig. 9). Starting with H1 = H3 = 0 and inactive A, H1 accumulation drives the system to the S state (first latch). However, it cannot return to G1 directly via H3, because without active B, the H3–A module is trapped in G2. Increasing B to 1 shifts the H3 nullcline downward and the A nullcline leftward, removing the G2 state and enabling transition to Meta (third latch). Returning to G1 requires B inactivation. An equivalent pair of B–H2 and B–H4 planes can be drawn, but they are fully analogous to Fig. 8 and therefore omitted.

**Fig. 9.**
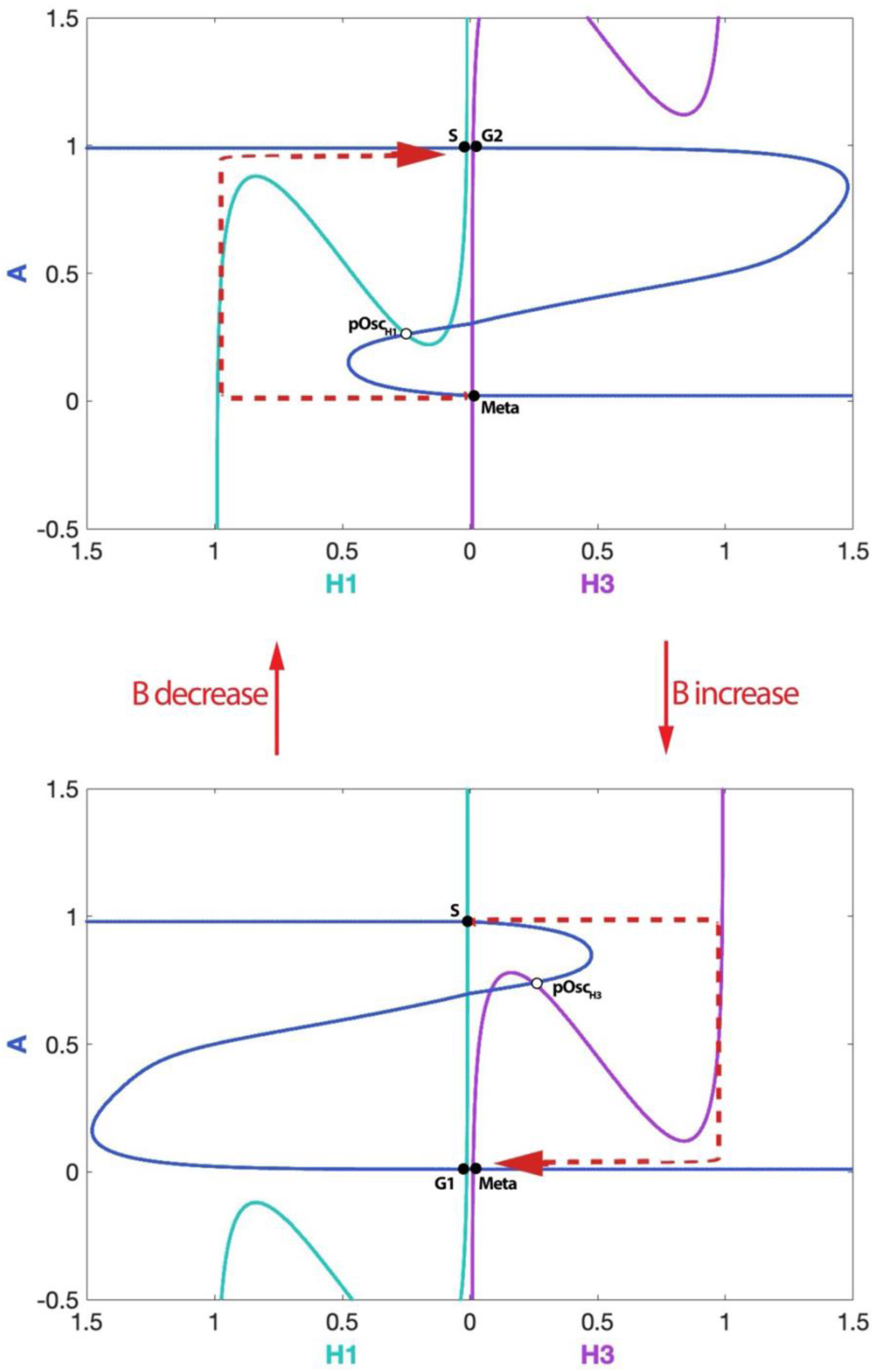
Latching gate view of the transitions in the core variable. **A** – In the top panel, the latching gate perspective shows that, when the core variable B is inactive, the H1-A system drives the G1/S transition. From the perspective of A, the state reached is identical to G2. Nevertheless, the cycle cannot be completed, unless B accumulates and the G2 state loses stability. As shown in the bottom panel, the H3-A system can complete bring the system to the Meta state, which from the perspective of A is identical to G1. The cycle is complete once B is eliminated.

## Discussion

### Summary

Progression through the cell cycle is governed by a complex molecular network [24] whose dynamics can include endo-oscillations. This category of phenomena has received considerable theoretical interest, such that several mechanistic and minimal models have been put forward. The present work shows that two previously proposed frameworks (Latching Gate and Newton’s Cradle) share a regulatory architecture consisting of two mutually regulated oscillators, despite starting from different assumptions. Importantly, this investigation highlights the link to previous models emphasising the role of multiple oscillatory units in cell cycle organisation [25] and particularly, in endocycle emergence [8,14]. This synthesis provides a framework for unifying major theoretical approaches to endocycles and clarifies the dynamical mechanisms underlying diverse cell cycle phenomena.

The double-oscillator concept is generalised to a four-oscillator architecture, with each oscillator driving a distinct cell cycle transition. In this framework, the cycle consists of stable pseudo-steady states separated by high-energy barriers; over longer timescales, each state generates a “force” through oscillatory modules to cross the next barrier. Endocycles arise when one or more barriers are removed, enabling spontaneous transitions without engaging certain helper modules.

### The tetra-stable system and physiological endocycles

A key property of this system is that any of the four states – or combinations thereof – can be suppressed by parameter changes. This flexibility accommodates all known physiological endocycle types. Endoreplication corresponds to loss of G2 and Meta, allowing repeated DNA replication without mitosis. Endomitosis retains G2 but suppresses Meta, so mitotic entry occurs but division fails – consistent with recent evidence that endomitosis might emerge from true cell cycle control network perturbations rather than mere cytokinesis defects [13]. Mitotic endocycles can be explained analogously: if G1 and S are absent but G2 and Meta remain, the cell enters mitosis multiple times without intervening DNA replication (e.g., meiosis, reductive mitosis). In some mitotic endocycles, mitotic entry recurs without replication or division (e.g. Cdc14 [8] and Cdc20 endocycles [9,11]) suggesting only the G2 state is stable.

The present dynamical definition of endocycle is very broad and many endocycles possible within this framework have no physiological correspondent (e.g. Fig. 8B,C). However, this broader notion can be applied to a range of cellular phenomena which have previously not been recognised as endocycles, but share common features. Under-replication [26] fits a case where S and G2 are suppressed, so cells move from G1 to M too quickly for complete DNA synthesis. Similarly, cells failing to enter quiescence after division under strong mitogen stimulation [27] resemble an endocycle with G1 suppressed. Thus, the model explains a broad range of phenomena using a single mechanistic principle.

The dynamics of endocycle emergence differ markedly between the two-state and four-state models, leading to a testable prediction. In the two-state model, and in the previously published Newton’s Cradle system, endocycles arise when one oscillator arrests – via a global SNIC or homoclinic bifurcation – producing critical slowing and an abrupt drop in oscillation amplitude for the disengaged variable. In the four-state model, by contrast, endocycles occur not by oscillator arrest but by minimising time in a helper- activating state, causing a gradual amplitude decline with increasing perturbation strength. This distinction can be probed experimentally: if perturbations (e.g., inhibitor titration) are applied incrementally, a sharp transition with critical slowing would favour a Latching Gate/Newton’s Cradle-like mechanism, whereas a gradual shift would be more consistent with the four-state architecture.

### The tetra-stable system and checkpoints

In this framework, checkpoints stabilise a cell cycle phase even in the presence of its corresponding helper, positioning checkpoints and endocycles as opposite extremes of phase duration control. A mechanistic model of the budding yeast cell cycle [23] proposed that mitotic oscillations arise from double-amplified negative feedback between Clb and Cln cyclin dependent kinases, producing a Clb–Cln phase plane akin to the tetra-stable core in Fig. 4B, but containing only the pOsc state. As a result, the system oscillates freely. In that model, checkpoint activation altered the nullclines to introduce stable steady states, arresting oscillations – an effect opposite to that of helper variables in the present framework. This hints at a potential mapping of the species in the abstract framework to cell cycle regulators.

### A molecular mapping of the tetra-stable system

In mammalian cells, variables A and B correspond to Cyclin A and Cyclin B, respectively. This mapping clarifies the meaning of the G1, S, G2, and M states, which differ in cyclin concentrations (and associated CDK activities) [28]. These regulators engage in multiple positive and double-negative feedback loops, generating bistability [29]. CycA:CDK2 stimulates CycB expression and inhibits its antagonists (e.g., APC/C:Cdh1, p21). Even though CycB:CDK1 can also promote CycA accumulation by targeting the same inhibitors, it suppresses CycA at key stages, stimulating its degradation in early mitosis and potentially downregulating its expression in S and G2. Such interactions may underlie the negative feedback between A and B assumed in the model.

The four helpers correspond to proteins that modulate cyclin levels. H1 aligns with Cyclin D [30], which initiates cyclin transcription ahead of G1/S, but declines with S-phase entry due to degradation [31]. H2 maps to the transcription factor FOXM1, which is activated by CycA:CDK2 [32] (and CycB:CDK1 [33]) and stimulates CycB expression [34]. H3 and H4 represent two functional states of APC/C:Cdc20, which sequentially target CycA and CycB for degradation during mitosis. APC/C activation by CycB:CDK1 [35–37] initially degrades only CycA due to partial inhibition by the mitotic checkpoint complex (MCC) [38]. Once the Meta state is reached and the mitotic checkpoint is silenced, APC/C:Cdc20 also degrades CycB [39]. This mapping is summarised in Fig. 10 and expanded in Appendix B.

**Fig. 10.**
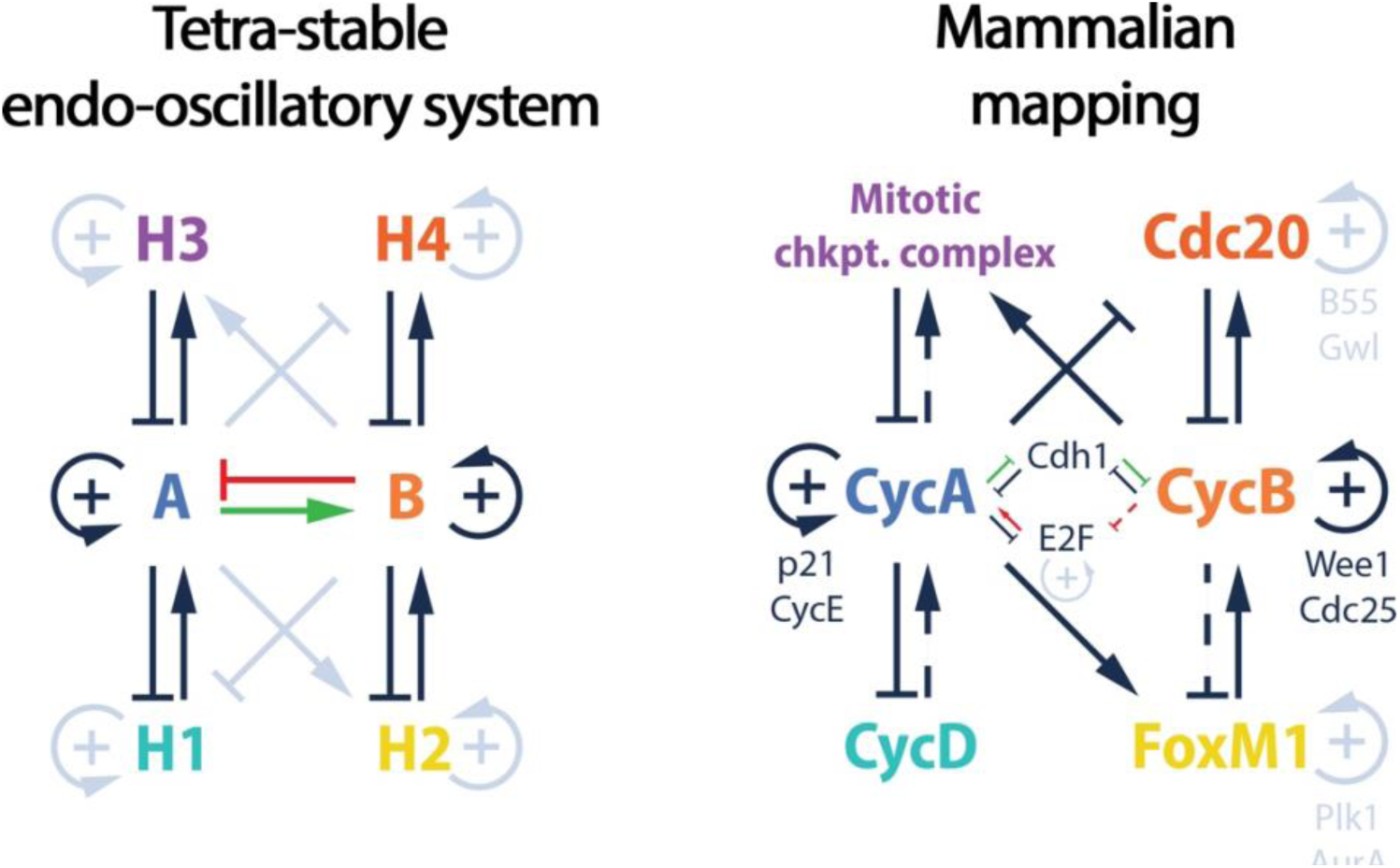
Mapping of the tetra-stable endo-oscillatory network to the mammalian cell cycle control network.

### The Wheel of Fortune analogy

The model can be visualised as a “wheel of fortune” with four equal sectors, each representing a cell cycle phase (Fig. 11). Between sectors are pegs that block the wheel’s motion unless the indicator’s spring is pulled with sufficient force. Spinning the wheel counterclockwise moves the indicator through the phases, but inertia alone cannot sustain motion—the pegs must be overcome repeatedly.

**Fig. 11.**
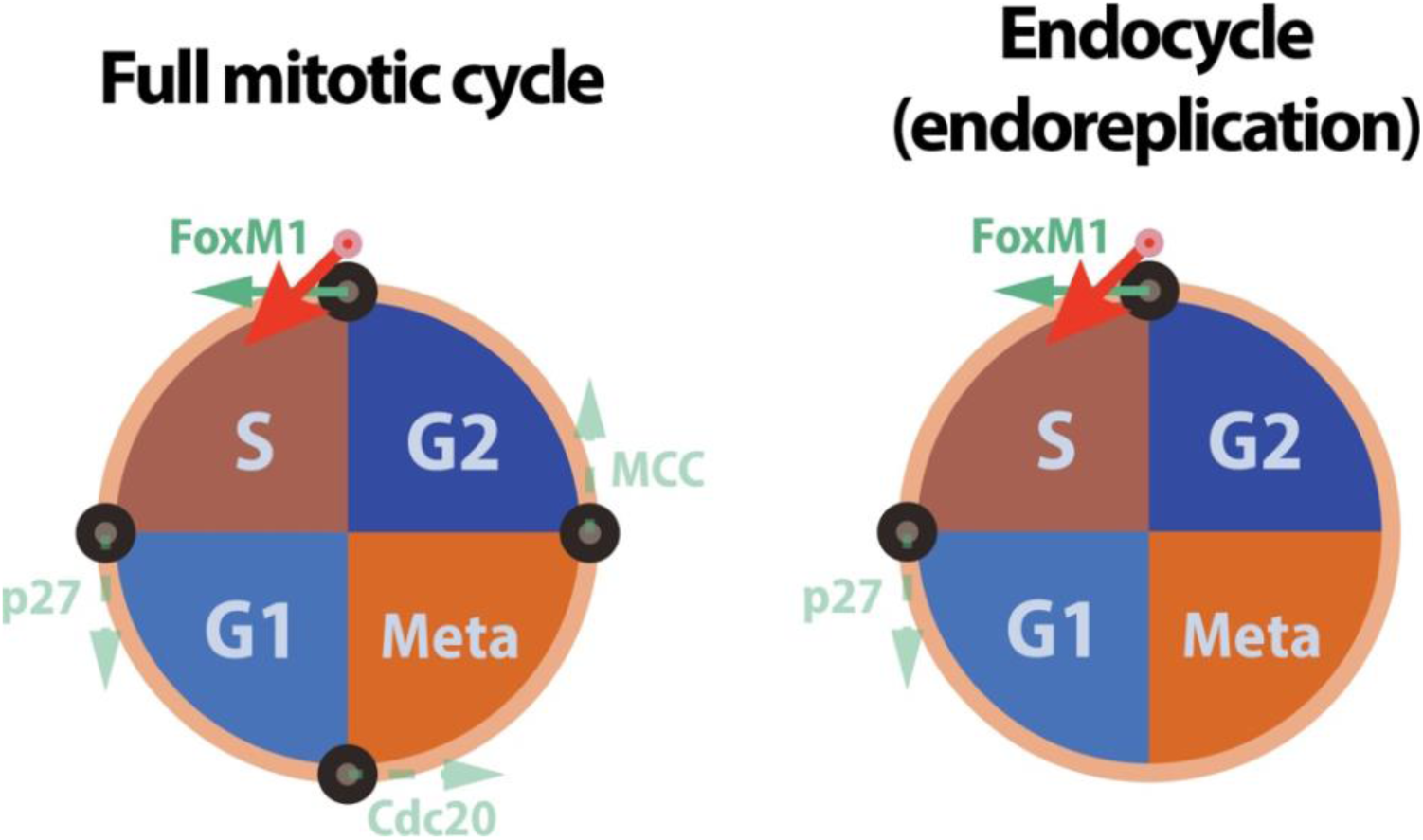
The Wheel of Fortune analogy.

In the cell cycle, this driving “force” comes from the helper variables, which promote cyclin synthesis or degradation to push the system into the next phase. An endocycle occurs when one or more pegs is removed, allowing the wheel to pass freely through the corresponding sectors without additional force.

Conversely, a checkpoint functions like a counterforce, resisting the helpers’ push and delaying phase progression. The CycA–CycB amplified negative feedback provides the intrinsic oscillatory potential, analogous to a frictionless wheel without pegs that can spin indefinitely. The “pegs” represent the energy barriers to transitions, making helper-driven forcing essential for regular cell cycle progression.

### Outlook

In essence, this paper has sought to clarify the conceptual landscape of endo-oscillatory phenomena in the cell cycle. Future work will aim to develop rigorous mechanistic models of the pathways outlined here and to test their predictions in vivo. More broadly, the principles uncovered in this study may extend to other physiological contexts where endo-oscillations are likely to occur – from ensembles of neurons, to juxtacrine signalling during development [40], to the coordination of periodic processes by the circadian clock.

## Materials and methods

### The four-oscillator model

The core system consists of two dynamical variable, A and B, defined according to 2 ODEs. Each equation describes the symmetric conversion of each species between two forms: an active one (e.g. A), and an inactive one (e.g. 1-A).

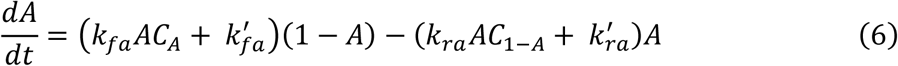

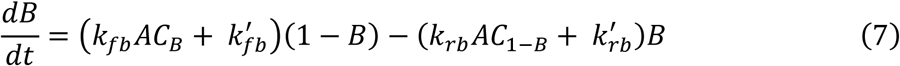

Both the positive and the negative terms of each equation contain a non-linear ‘auto- catalytic’ (AC) function, which provides the coupling of the two variables. The form of these expressions is that of a Hill function whose threshold of activation depends on the opposite dynamical variable. Thus, A stimulates its own activation, but is opposed by B, as the threshold of the Hill function increases linearly with B:

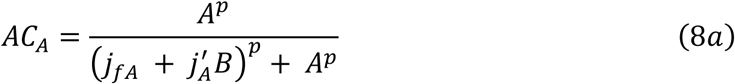

Similarly, the inactive form of A, denoted by 1 – A, is also self-stimulated, but opposed by 1 – B (i.e. stimulated by B).

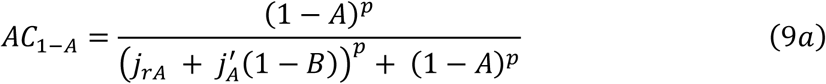

Thus, B inhibits A through both terms. This description is necessary for rendering the dynamical system fully symmetrical and greatly simplifies the model further down the line. The auto-catalytic functions of B are treated similarly, but note that A stimulates the conversion of B to the active form:

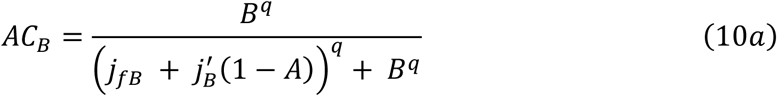

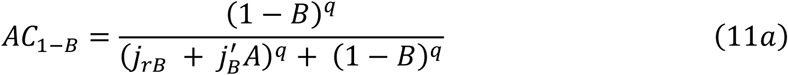

The helper variable H1 is modelled in analogy to A and B, such that H1 is present in two forms: active (H1) and inactive (1 – H1). The conversion between the two forms is done basally (rate constants 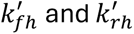) and autocatalytically. In addition, a checkpoint term is present, which favours H1 conversion to the inactive form with mass action kinetics. C1 is a parameter taking values between 0 and 1, and 𝑘_𝑐_ is a kinetic parameter.

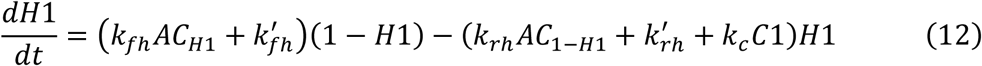

The autocatalytic terms show that the activation of H1 is inhibited by both A and B, as H1 accumulation is triggered when both A and B are inactive:

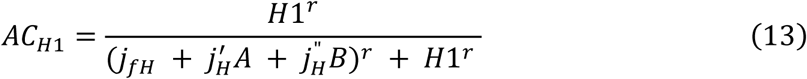

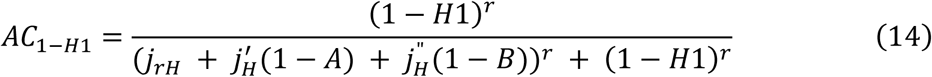

All other helpers are modelled analogously, except A and B terms are replaced by (1 – A) and (1 – B), and vice versa, if A or B activate the helper instead of inhibiting it. For example, H2 is activated by A and inhibited by B. Thus, its auto-catalytic functions are modelled as (changes in bold):

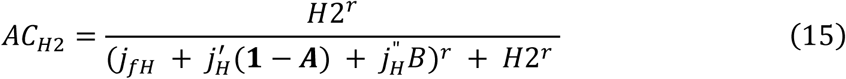

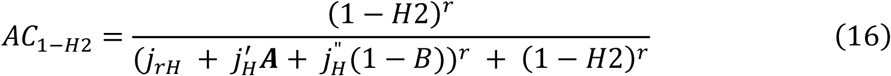

To link the helpers to the core variables, the core’s auto-catalytic functions are updated to:

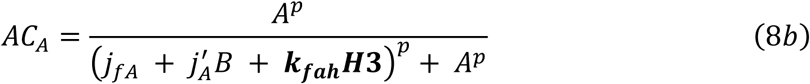

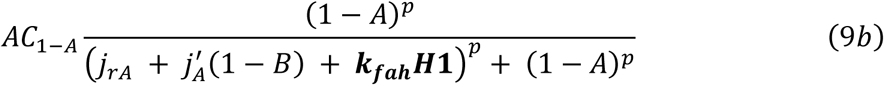

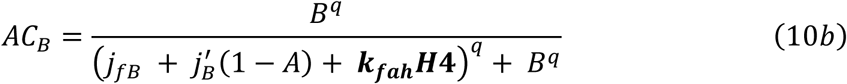

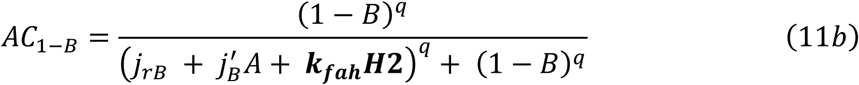

As such, H1 activates A and H3 inactivates it. Similarly, H2 activates B and H3 inactivates it.

### Parameter values

#### Computation

The models were implemented in the freely available software XPPAUT (https://sites.pitt.edu/∼phase/bard/bardware/xpp/xpp.html). The code for .ode files used to simulate the models and calculate phase planes and bifurcation diagrams are provided in Supplementary materials. The .ode files were used to plot figures as per Table 3.

**Table 3.**
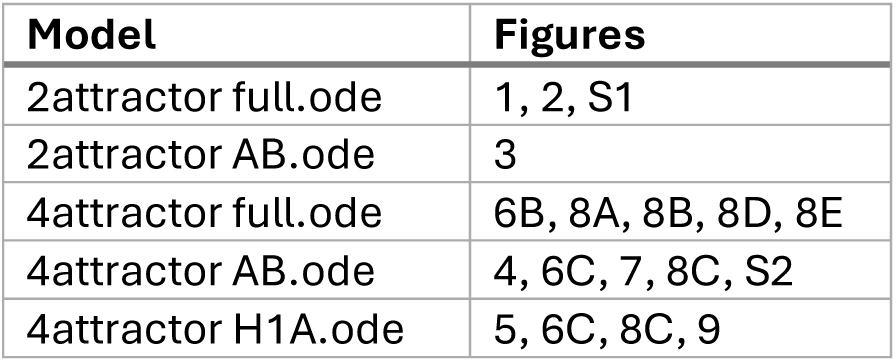
The .ode files and the figures for which they were used.

| **Model** | **Figures** |
| --- | --- |
| 2attractor full.ode | 1, 2, S1 |
| 2attractor AB.ode | 3 |
| 4attractor full.ode | 6B, 8A, 8B, 8D, 8E |
| 4attractor AB.ode | 4, 6C, 7, 8C, S2 |
| 4attractor H1A.ode | 5, 6C, 8C, 9 |

## Acknowledgements

I thank Bela Novak and John Tyson for useful discussions and the critical reading of the manuscript. This work was funded by the Elizabeth Murphy Scholarship, awarded by Trinity College, Oxford.

### Appendices

#### Appendix A: Justification for Latching Gate modelling approach

Although the variables A, H1 and H2 in the model do not correspond one-to-one with specific cell cycle regulators, they can be related phenomenologically to known biochemical activities. Variable A represents the pooled CDK activity (S-CDK and M- CDK), with low A corresponding to G1 and high A corresponding to S/G2/M. CDKs participate in multiple positive feedback circuits (both double positive and double-negative feedback). For example, this is achieved by phosphorylating and inactivating negative regulators, such as the APC/C adaptor Cdh1 (active in G1), stoichiometric inhibitors such as p21, and tyrosine kinases such as Wee1, or activating positive regulators, such as the Cdc25 phosphatase.

The functions B and B′ mediate these positive feedback effects. They are represented as steady-state expressions rather than explicit ODEs because protein phosphorylation typically occurs on much shorter timescales than cyclin synthesis or degradation, which is usually the rate-limiting step in CDK regulation. Many CDK targets show sigmoidal responses to CDK activity due to ordered, distributive multisite phosphorylation [41,42], a property captured here with Hill functions. The combination of strong positive (or double-negative) feedback and nonlinearity is well established as a mechanism for bistability in cell cycle regulation, as assumed for A.

The helpers H1 and H2 represent proteins that provide negative feedback on CDK activity. In mammalian cells, such feedback can be mediated by E2F transcription factors or the APC/C adaptor Cdc20 [11]. In the model, H1 and H2 act on the placeholder functions B and B′ for mathematical convenience. However, in vivo, negative feedback more often acts directly on cyclin synthesis or degradation. As for the biochemical analogues of B and B’, the helpers respond nonlinearly to CDK thanks to ordered, distributive multi-site phosphorylation. However, because the helpers are modelled as dynamic variables, rather than steady state expressions, it is not convenient to use a Hill function directly, hence why Michaelian kinetics were used to describe the conversion between the active and inactive forms.

#### Appendix B: Molecular Mapping – Additional Mechanistic Details

The interplay between helpers and cyclin levels can be more complicated than the idealised model might suggest. This is expected, given the simplicity of the toy framework relative to the underlying biological network. Nevertheless, it can be argued that the biochemical control mechanisms still follow the logic outlined in the Discussion.

Although Cyclin D does not upregulate Cyclin A directly, it partially inhibits the transcriptional repressor Rb, leading to E2F activation and Cyclin E accumulation. Cyclin E:CDK2 then initiates a positive feedback loop that inactivates APC/C:Cdh1, allowing Cyclin A accumulation and S-phase entry [44]. Unlike H1, Cyclin D reaccumulates before cell division [28], but unscheduled Cyclin A build-up is avoided through multiple mechanisms, including E2F inhibition via phosphorylation and Cyclin A degradation during mitosis.

CycB:CDK1 does not directly inhibit FOXM1, as B does H2 in the model. However, CDK1 activity in M-phase triggers chromosome condensation, which shuts down transcription [45], indirectly suppresses Cyclin B. This reasoning also applies to Cyclin A, providing an alternative pathway for the Cyclin A - Cyclin B negative feedback discussed in the main text.

The mitotic checkpoint complex (MCC) alone is an insufficient analogue for H3, since CycA does not directly activate MCC. Instead, H3 is better viewed as a composite of MCC with upstream mitotic regulators (Wee1, Cdc25, CDK1). Both CycA [46] and CycB [47,48] activate Cdc25 and inhibit Wee1, leading to CycB:CDK1 activation. Activated CDK1 phosphorylates APC/C [35–37], enabling MCC formation.

The mapping of H4 to APC/C:Cdc20 is more direct. CycB:CDK1 activates Cdc20, whereas CycA:CDK2 inhibits it [49]. As previously noted, APC/C:Cdc20 targets CycB for degradation directly during anaphase.

In summary, the mapping between the dynamical variables and molecular species reflects either single proteins or functional modules within the cell cycle control network. While simplified, this mapping captures core regulatory logic that may underlie the diverse endo-oscillatory behaviours explored in this study, even if the full physiological system contains additional layers of complexity and cross-talk.

## Supplementary figures

**Fig. S1.**
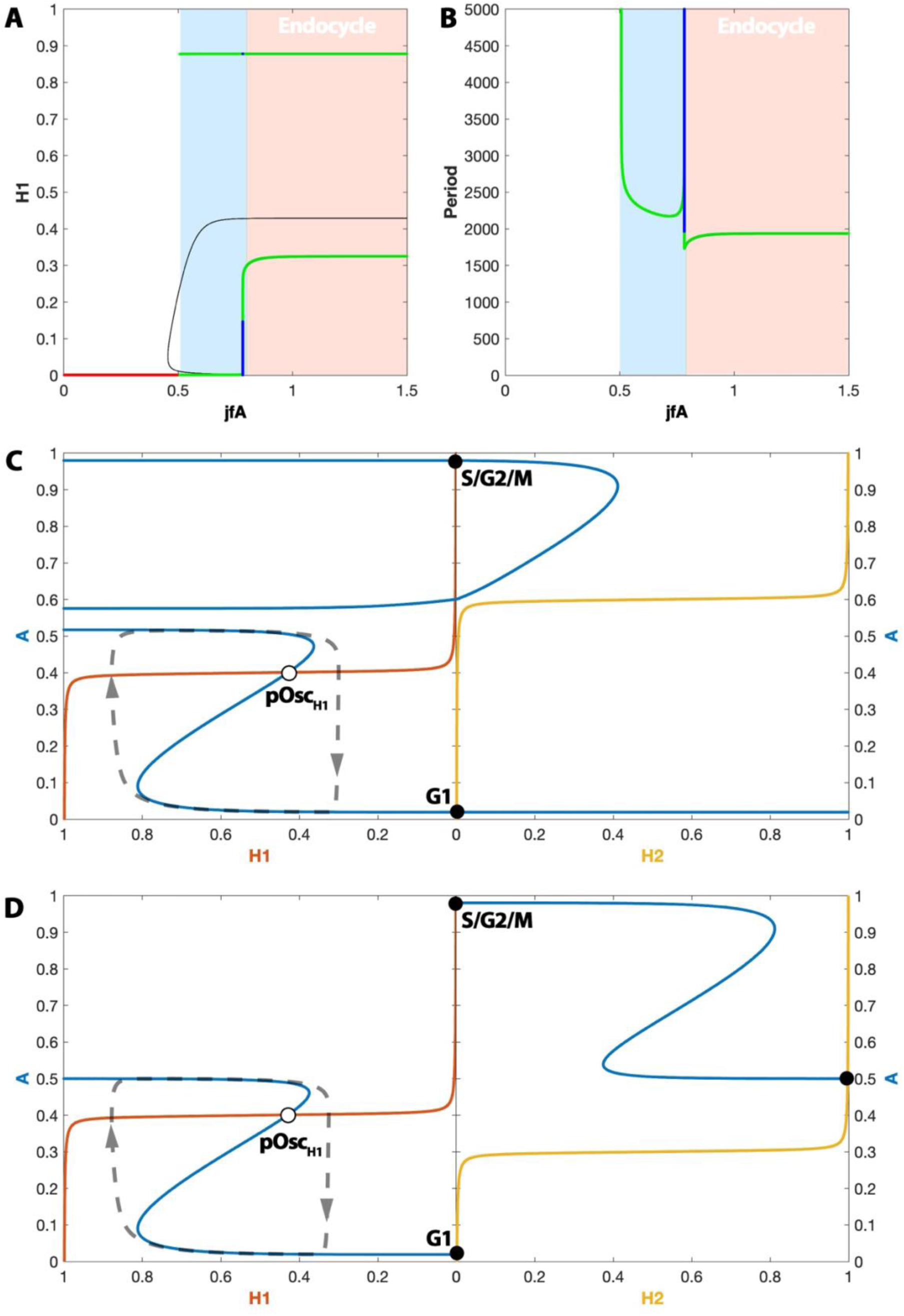
Endocycle dynamics in the minimal Latching Gates model. – (A) Bifurcation diagram of H1 with respect to the parameter *jfA*. The plot shows that even though the amplitude of H1 is reduced sharply for *jfA* values above the threshold for endocycle induction, unlike H2, H1 still has a substantial contribution to the oscillation. Green indicates the maximal and minimal amplitude of stable limit cycle solutions, while blue indicates the amplitude of unstable limit cycles. (B) Plot of oscillation period as a function of the *jfA* parameter. At the boundary between the ‘complete’ mitotic oscillation and the endocycle regime, the period of the unstable oscillation approaches infinity at a very fast rate. This is consistent with critical slowing down at the value of *jfA* for which a limit cycle oscillation emerges/disappears at a homoclinic bifurcation. (C) The A-H1 phase plane with H2 = 0 and A-H2 phase plane with H1 = 0, assuming *jfA* = 0.8. The plot illustrates that the H1 endocycle can emerge if the pOsc_H2_ state is suppressed, even when the S/G2/M stable steady state continues to exist. (D) The A-H1 phase plane with H2 = 1 and A-H2 phase plane with H1 = 1, assuming *kfh2* = 0.7.

**Fig. S2.**
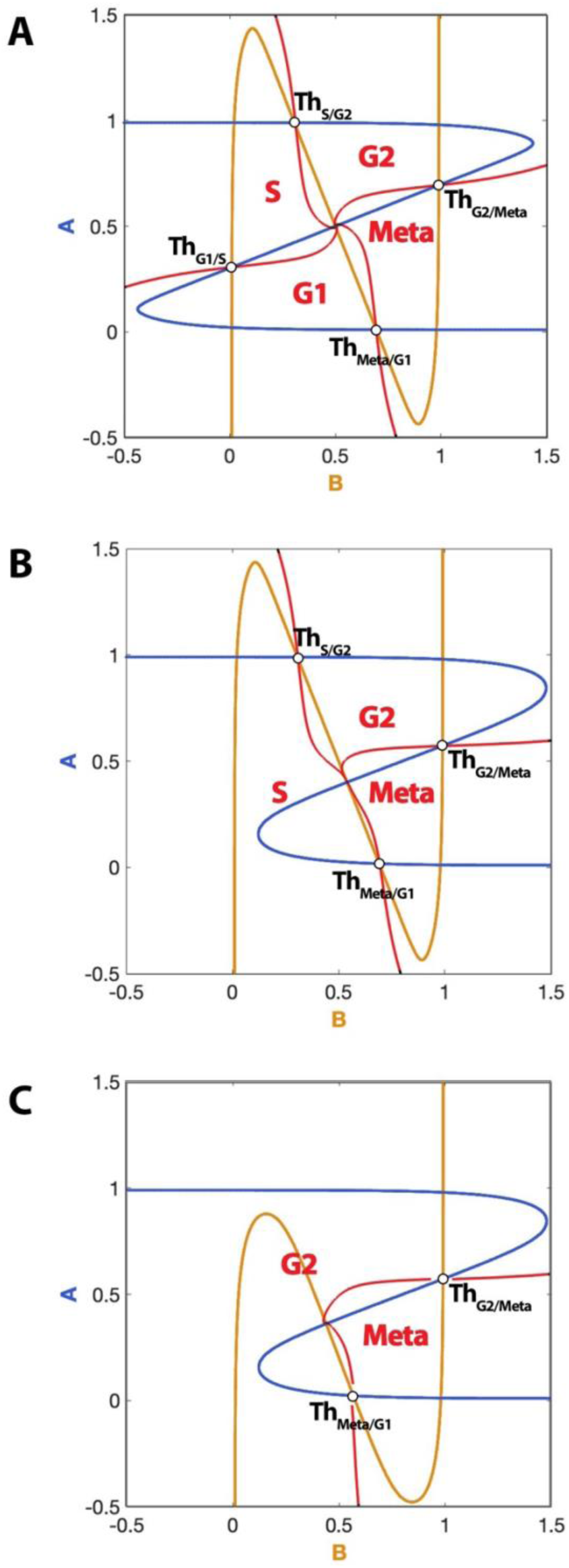
The basins of attraction of the stable steady states in the tetra-stable core model. The boundaries of the basins of attraction are drawn is red for (A) the basal system, (B) the system perturbed with *jfA*=0.8 (G1 suppressed) and (C) the system perturbed with *jrA*=*jrB*=0.8 (G1 and S suppressed).

## XPPAUT .ode scripts

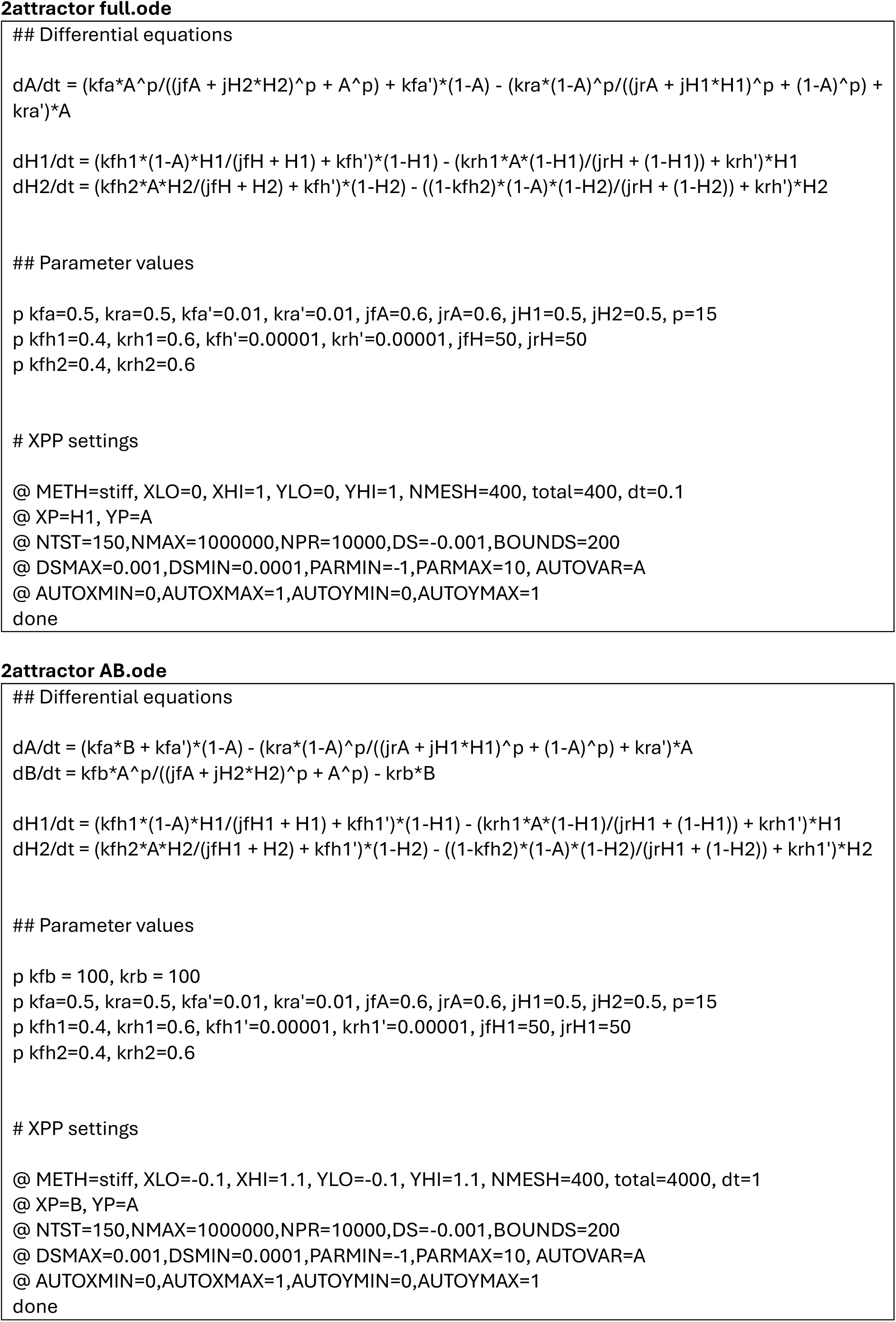
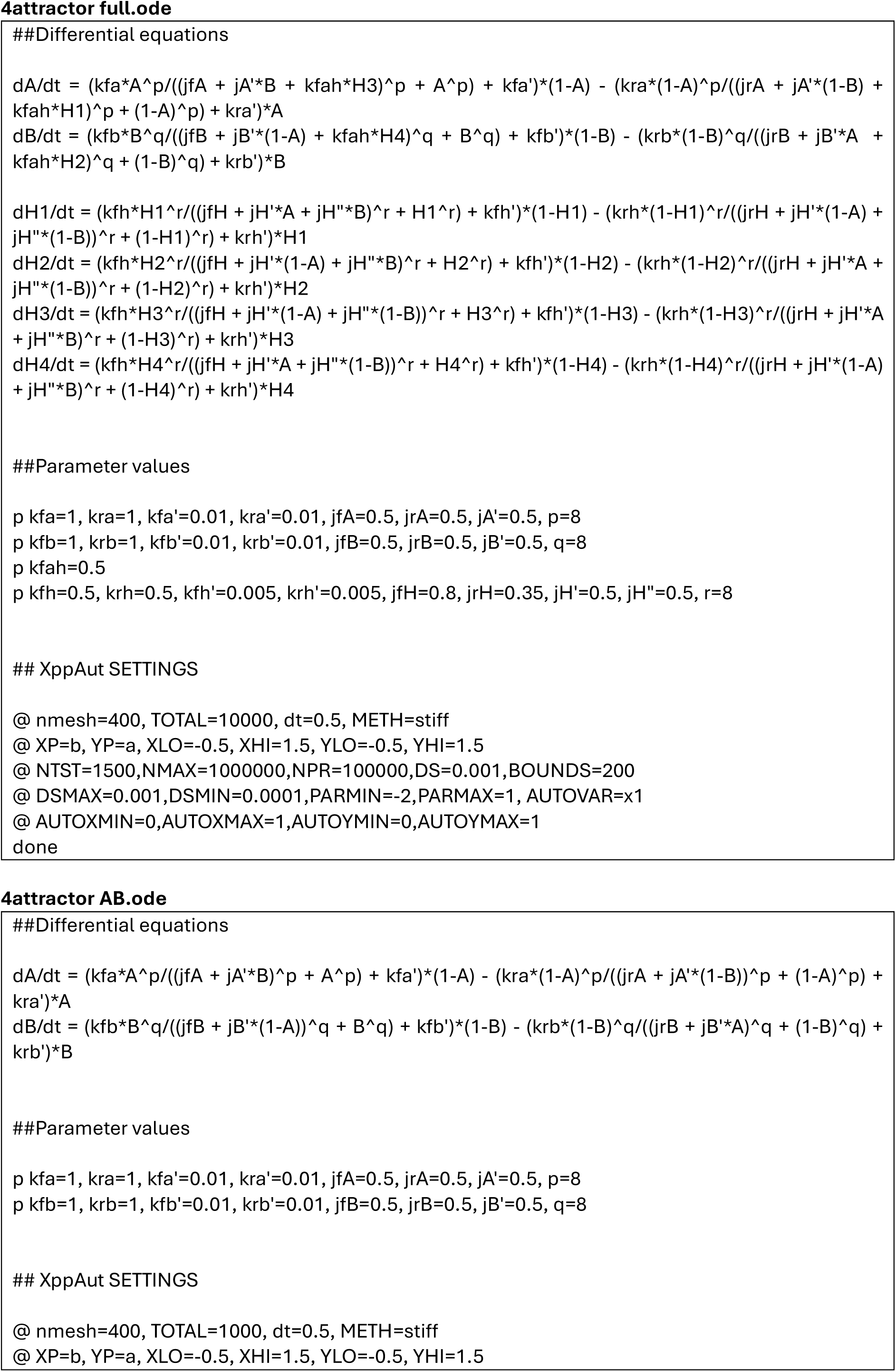
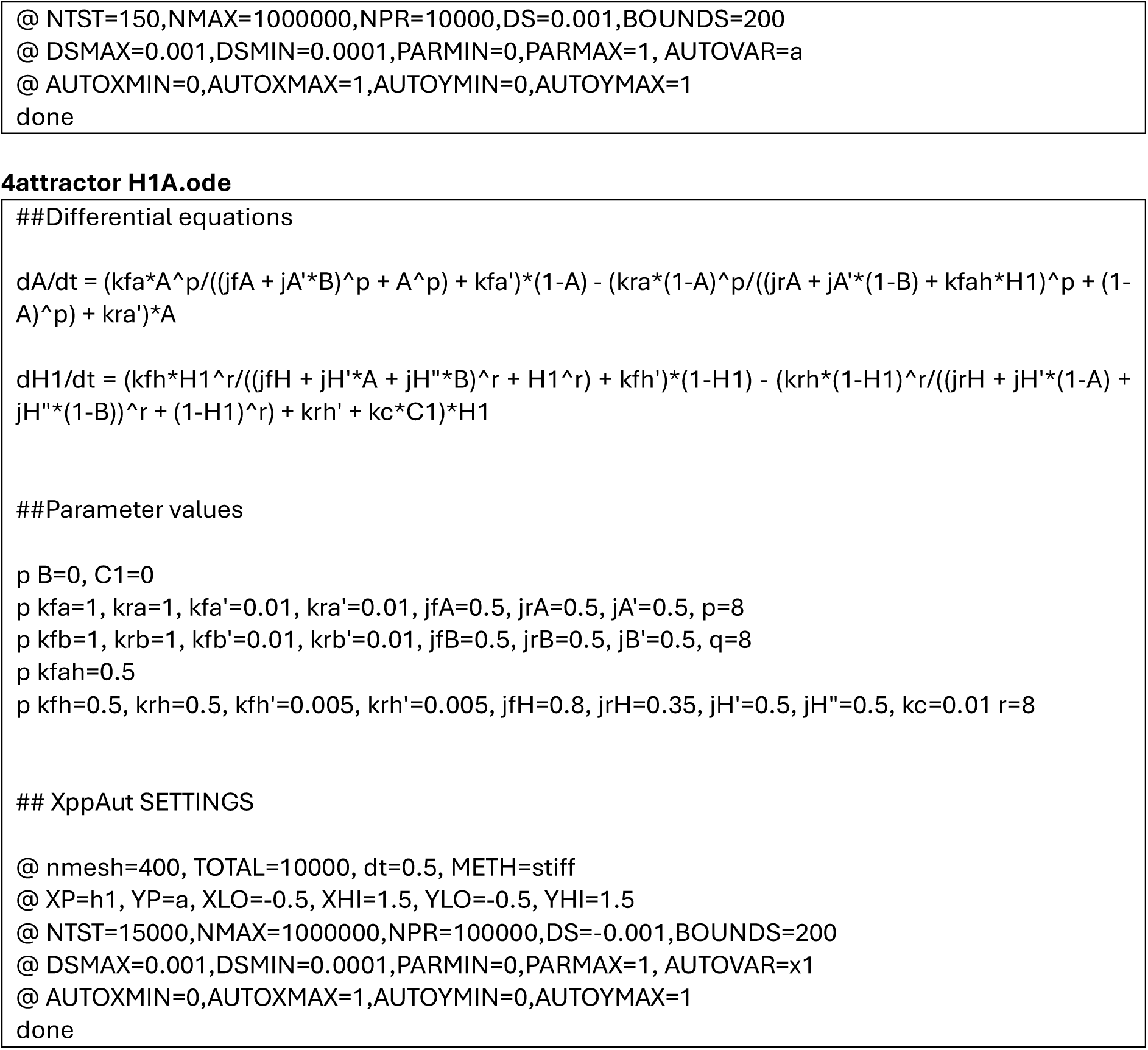

